# Changes in the translational landscape during red clover necrotic mosaic virus infection

**DOI:** 10.64898/2026.06.20.733525

**Authors:** Pulkit Kanodia, Zachary Lozier, Filip Lastovka, David Walker, Peng Liu, Betty Chung, W. Allen Miller

## Abstract

Viruses alter host gene expression to create a proviral environment while the host simultaneously regulates gene expression to restrict the virus spread. Owing to RNA virus’ complete reliance on the host translational machinery, it is important to assess the translational control during virus infection. Therefore, we used ribosome profiling (ribo-seq) paired with RNAseq to observe how red clover necrotic mosaic virus (RCNMV) infection of Arabidopsis plants alters cellular gene expression at the levels of mRNA abundance and translation efficiency. We determined that at 5 days post-inoculation (dpi), the translational response to RCNMV infection is enriched in genes of the innate immune system. Expression of a tumor necrosis factor receptor-associated factor (TRAF)-like protein, a regulator of development and immune response, was translationally but not transcriptionally upregulated early in systemic infection. By 8 dpi, many pathways were regulated/dysregulated, and unfolded protein response (UPR) genes were transcriptionally upregulated but with reduced translation efficiency. Ribosome profiling of RCNMV RNAs revealed (i) -1 programmed ribosomal frameshifting at 7.5-8.0%, the first direct measurement of frameshift efficiency in infected cells for any plant virus; (ii) that coat protein is translated at extremely high efficiency, while the RNA-dependent RNA polymerase is translated least efficiently, and (iii) an unexpected extremely strong ribosomal pause site in the open reading frame that encodes the movement protein. To our knowledge, this is the first genome-wide study that assesses the translational control of gene expression in plants infected with a virus from the large and diverse *Tombusviridae* family.

**Importance:** Positive strand RNA viruses usurp the host’s translation machinery to synthesize viral proteins. Moreover, translation of host mRNAs is altered by virus infection, both as part of the host immune response and by the virus to inhibit host defenses. To assess all these changes globally, we used ribosome profiling of plants infected with a member of the large and ubiquitous Tombusviridae family. We identified key host genes and pathways that were differentially altered in translation efficiency, giving us an understanding of host responses not detectable by conventional RNA sequencing. Moreover, ribosome profiling revealed (i) the most accurate calculation of efficiency of ribosomal frameshifting during infection for any plant virus, (ii) the extremely high level of translation of viral coat protein, and (iii) an unexpected strong ribosomal pause site in the movement protein gene. This work provides understanding of a new dimension of gene expression control in plant-virus interactions.

## Introduction

Plant viruses induce changes in host gene expression to create a pro-viral cellular environment, while the host regulates the cellular and viral gene expression to restrict virus infection (1–4). Understanding host-virus interactions on a genome-wide scale is valuable for identification of the pro- and antiviral pathways that can be exploited and targeted to develop virus-resistant plants (5–7). While numerous studies have elucidated the transcriptomic and post-transcriptional regulation of cellular and viral gene expression (8–10), only a few studies have assessed host translation regulation in plant viral infection (11–15).

Previously, high-throughput sequencing of polysome-associated mRNAs has been used to study translational control of plant mRNAs during pathogen infection (16–19) but this technique can suffer from limited resolution and accuracy (20). In contrast, ribosome profiling (ribo-seq) has enabled researchers to study translational control of gene expression in a quantitative, high-throughput manner at a genome-wide scale with single-nucleotide resolution (20). Ribo-seq is a variation of RNA-seq in which translating ribosomes are digested with ribonuclease to degrade all mRNA not protected by the 80S ribosome. The resulting 28-30 nt ribosome-protected fragments (RPFs) are deep sequenced. The number of RPF reads mapping to each mRNA is proportional to the number of ribosomes translating that mRNA. RNA-seq of the same samples is used to determine changes in mRNA levels to allow calculation of changes in translational efficiency (TE, RPFs per mRNA).

Ribo-seq has been applied to explore the translation landscape of the cellular and viral mRNAs during infection in animal systems (21–27) or the translatome in plants in absence of viruses (28–30) but there are only a few published studies that apply ribo-seq to plant virus-host interactions (31–35), with most of these focusing only on the viral translatome. None of the studies focused on plants infected by a member of the *Tombusviridae*, which is of particular interest, as these viruses harbor cap-independent translation elements that bind translation initiation factors. Thus, infection by these viruses has potential to impact host translation. To investigate the effect of tombusvirid infection on host translation, as well as to characterize translation of the viral genome, we used *Arabidopsis thaliana* and red clover necrotic mosaic virus (RCNMV) as a model system.

RCNMV (genus *Dianthovirus*, family *Tombusviridae*) is a bipartite RNA virus with uncapped and non-polyadenylated positive-sense single-stranded genomic RNAs 1 and 2 (36, 37). Replication protein, P27, and a -1 programmed ribosomal frameshift product, P88, that contains the RNA-dependent RNA-polymerase (RdRp) domain, are translated directly from RNA1 (37–39). Coat protein (CP) is translated from a subgenomic RNA (CPsgRNA1) derived from the 3’-terminal one-third of RNA1. CP is required for long-distance movement in the plant (40, 41). Movement protein (MP) is translated from RNA2 and is required for cell-to-cell movement (40, 42). Additionally, a noncoding subgenomic (ncsg) RNA, called SR1f, comprising most of the 3’ untranslated region (UTR) of RNA1 accumulates in infected cells. SR1f is generated as a stable degradation product formed by incomplete degradation of RNA1 and CPsgRNA1 by the host 5’ to 3’ exoribonuclease (43, 44). Translation of viral proteins from RNA1 and CPsgRNA1 occurs via a 3’ cap-independent translation element (CITE) in the 3’ UTR, called TE-DR1 (45). TE-DR1 belongs to the barley yellow dwarf virus-like translation element (BTE) class of CITEs that recruit the translation machinery via binding to translation initiation factor eIF4G (45–48). TE-DR1 is present also in the 3’ co-terminal SR1f, thus we hypothesize that SR1f may sequester the host translation machinery and spatio-temporally regulate viral as well as cellular translation (14, 43). In addition, the tombusvirid tomato bushy stunt virus (TBSV) replicase complex employs key proteins involved in translation such as translation elongation factor eEF1A, DEAD-box helicase DDX3, and hsp70 (49–52). All of the above interactions may affect host translation in unpredictable ways.

To investigate how RCNMV infection in Arabidopsis affects cellular gene expression, both at the level of transcription and translation, we performed ribosome profiling (ribo-seq) of plants at early and late stages of systemic infection. We identified genes that are differentially regulated owing to the change in their (i) mRNA abundance without any change in the number of ribosomes translating those mRNAs, (ii) translation efficiency, i.e., the number of translating ribosomes without any change in the mRNA abundance, and (iii) mRNA abundance as well as translation efficiency (53). We found that the translationally regulated genes, only in the early phase of systemic infection, are enriched in protein domains that are involved in plant defense responses. In addition to the host gene expression, we also assessed the ribosome profiles on viral RNAs which displayed the translation of the four known ORFs in the RCNMV genome. Ribo-seq also allowed visualization of -1 programmed ribosomal frameshifting for the translation of p88 (RdRp) ORF, and revealed an unexpected, extremely strong pause site in translation of RNA2. As expected, we found out that among the four ORFs, p88 ORF is translated least efficiently, and the CP ORF is translated most efficiently.

## Results

### RCNMV accumulation increases steeply at 6 to 7 dpi in DCL2/4-knockout Arabidopsis

We tested Arabidopsis as a host because of its well annotated, small genome. However, RCNMV infection in wildtype Arabidopsis Col-0 was asymptomatic and very uneven (54). Thus, we reduced the innate immune response by using DCL knockout mutants (dcl2-1/dcl4-2t), which lack DCL2- and DCL4-dependent production of virus-derived siRNAs, while leaving canonical pathogen-triggered immunity, salicylic acid signalling, and the unfolded protein response genetically unaltered. Among the four DCL proteins (DCL1, 2, 3, 4) in Arabidopsis, DCL2 and DCL4 generate virus-derived siRNAs required for antiviral RNA silencing (55). Arabidopsis dcl2-1/dcl4-2t (56) is a loss-of-function double knock-out line with non-functional antiviral RNA silencing machinery. These knockout plants appear normal when not infected.

Our first objective was to determine an early and a late time-point at which the systemically infected leaves would be harvested for ribo-seq. Because change in translation is a quicker response than transcription, we speculated that at early time-points prior to symptom development, when virus levels are low, we might see more translationally regulated changes in gene expression. The late time-point refers to the timepoint after the appearance of symptoms with a high viral RNA accumulation. Changes in translation at this timepoint could be due to effects of accumulated viral proteins and RNA or due to signaling from infected to uninfected cells. A time course assay from 4 to 10 days post inoculation (dpi) demonstrated the appearance of strong symptoms at around 7 dpi (Supplemental Fig. S1A). Symptoms included chlorosis, necrosis, and severely mosaic and epinastic leaves. Subsequently, RCNMV RNA1 accumulation was assessed using RT-PCR and qRT-PCR (Supplemental Fig. S1B, S1C). Consistent with the appearance of symptoms, viral RNA accumulation increased dramatically at 6-7 dpi in repeated experiments. Therefore, for ribo-seq, we chose 5 dpi as an early time-point when there are no symptoms, but RCNMV was present and replicating, and 8 dpi as a late time-point when all the infected plants were fully symptomatic and contained large amounts of virus but were not yet necrotic (Fig. 1).

**Figure 1.**
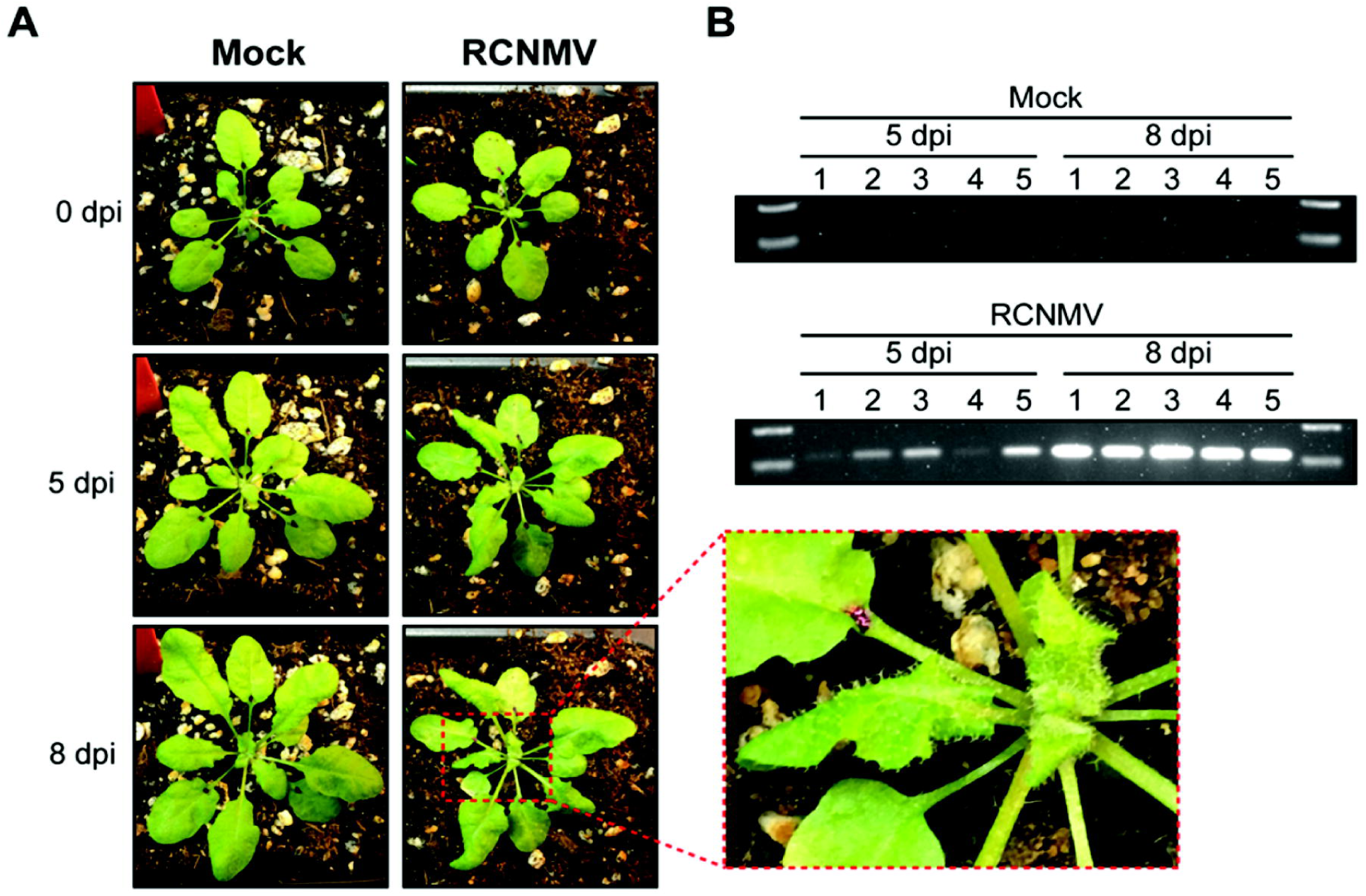
Plants used for ribo-seq and RNA-seq. (A) Symptoms at 0, 5, 8 dpi in mock- and RCNMV-infected samples. The zoomed-in image shows the symptoms on the non-inoculated systemic leaves. (B) RT-PCR to verify the accumulation of RNA1 in systemic leaves at 5 and 8 dpi in RCNMV-infected plants.

### Ribo-seq data show high triplet periodicity

For ribo-seq, initially we used five biological replicates for each mock and infected sample, however one of the mock-infected replicates was mis-handled, so we used only four biological replicates for the mock-infected amples (Supplemental File S1). In all samples, about half of the reads mapped to noncoding (mostly ribosomal) RNAs, which were removed from further analysis. Length distribution of remaining reads showed a dominant peak at 26–29 nt (Supplemental Fig. S2A), of which 11 million to 34 million reads mapped to the Arabidopsis genome, depending on the sample (Supplemental File S1 and Supplemental Fig. S3). No strong batch effects were detected by principal component analysis (Supplemental Fig. S4) and biological replicates were highly reproducible (Pearson’s r ≥ 0.92 for both RPF and RNA-seq counts) (Supplemental Fig. S4F, G).

To determine if the reads obtained from ribo-seq were true RPFs, we tested for triplet periodicity in the reads mapped to mRNAs. Triplet periodicity reflects the higher number of reads mapping to every third base in coding regions due to the ribosome pausing that occurs at each codon. The triplet periodicity for 26-29 nt ribo-seq reads and lack thereof, for RNA-seq reads, was obvious using metagene analysis (Ribotaper, Supplemental Fig. S2). Metagene analysis also showed that ribo-seq reads mapped predominantly to the CDS (Supplemental Fig. S2C). Furthermore, according to the number of nucleotides protected by the ribosomes upstream of the P-site (Supplemental Fig. S2C, see peaks upstream of start and stop codons), we can see that the 5’end of RPFs are preferentially digested by the RNase, as reported previously (57). The 5’ ends of RPFs are further upstream of the stop codon (14 nt for 28 nt RPFs) than the start codon (12 nt), owing to change in chape of the ribosome upon entry of release factor (Supplemental Fig. S2C). In contrast, the RNA-seq reads, obtained from the same lysates used for ribo-seq, mapped equally to both CDS and UTRs and showed no triplet periodicity (Supplemental Fig. S2C), as expected. All the above-mentioned data characteristics show the successful preparation of high-quality ribo-seq data where majority of the reads represent the true RPFs.

### Differentially expressed and translated genes

DESeq2 (58) was used for differential expression analysis separately for RNA-seq and ribo-seq data to identify differentially expressed genes (DEGs) and differentially translated genes (DTGs), respectively (All are listed in Supplemental File S2). We define DEGs as those having absolute (log_2_-fold change of RNA abundance) > 1 with adjusted p-values < 0.05, and DTGs as those having absolute (log_2_ fold change of RPF abundance) > 1 with adjusted p-values < 0.05. For DTGs, the change in RPF abundance can arise due to the change in mRNA abundance (DEGs), or change in translation efficiency, or both. Thus, DTG must not be confused with translational efficiency (TE). DTG simply represents a change in RPFs. This may result entirely from change in mRNA abundance, with no change in TE. In contrast, a change in TE requires that the change in RPF counts for a gene is not proportional to the change in mRNA abundance of that gene.

For RCNMV vs mock datasets at 5 dpi, we identified 356 DEGs and 500 DTGs whereas at 8 dpi we identified 3441 DEGs and 3263 DTGs (Fig. 2A). As expected, a far greater number of genes were differentially regulated at both the RNA and RPF levels at 8 dpi when all the plants were severely symptomatic. The scatter plot in Fig. 2B shows changes in mRNA level (DEGs) vs translation level (DTGs). The blue points are DTGs with non-significant change in mRNA abundance, green points are DEGs with non-significant change in RPF abundance, and red points are genes that are both DEGs and DTGs (Fig. 2B). For this scatter plot, gene IDs, gene product functions, and exact levels of change in mRNA and RPF levels are indicated in Supplemental file S2. To assess if the protein products from the translationally-regulated genes have any common features, we conducted protein-domain enrichment analysis using ThaleMine (59). For stringent criteria, we focused only on the DTGs (blue) in the yellow shaded area which are the most likely translationally regulated genes. Among the 101 upregulated genes at 5 dpi, several protein domains, such as protein kinases, leucine-rich repeat (LRR), Toll/interleukin-1 receptor homology (TIR), and Gnk2-homologous domains among others, were enriched (Fig. 2B). These domains are central to plant innate immunity (60–62). The six downregulated genes were insufficient for enrichment analysis. Even though there were far more translationally regulated genes at 8 dpi, no protein domains were enriched in the 280 upregulated and 250 downregulated genes (Fig. 2B).

**Figure 2.**
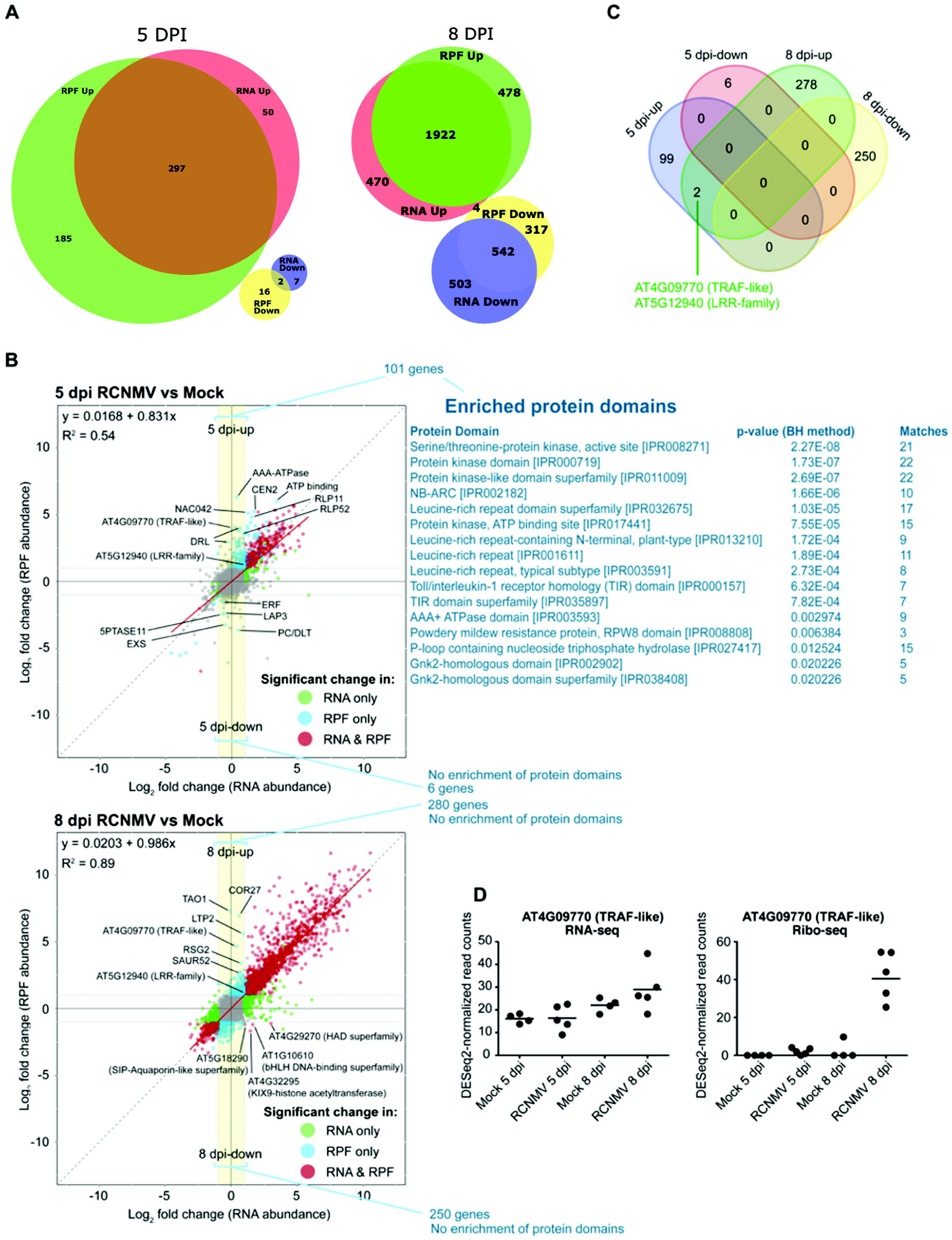
Differentially expressed and translated genes in RCNMV-infected plants. **(A)** Venn diagram showing the number of genes identified as DEG from RNA-seq data and DTGs from Ribo-seq data, **(B)** Scatterplot of log_2_ FC (RPF abundance) vs log_2_ FC (RNA abundance) with only the genes that had a valid (non-NA) DESeq2 output. **(C)** Venn diagram showing the number of translationally regulated genes at 5 and 8 dpi (blue-colored genes withing the yellow shaded region). **(D)** DESeq2-normalized read counts of AT4G09770 (TRAF-like) gene in RNA-seq and Ribo-seq datasets across all the treatments.

Among the translationally regulated genes (Fig. 2B, blue points in yellow shaded area), only two genes were common at both timepoints, namely AT4G09770 (TRAF-like family protein), that had a high change in RPF abundance, and AT5G12940 (LRR-family protein) (Fig. 2C), which had only a modest change in RPF abundance. By plotting the DESeq2-normalized counts for AT4G09770 (TRAF-like family protein) across ribo-seq and RNA-seq samples in all treatment conditions (Fig. 2D), we can see that this gene is actually upregulated at the level of translation. We also found four genes at 8 dpi, identified both as DEG and DTG, that were regulated in opposite direction, i.e., upregulated transcriptionally but downregulated translationally (Fig. 2B). These genes included AT5G18290 (SIP-Aquaporin-like superfamily protein), AT4G32295 (KIX9-histone acetyltransferase), AT1G10610 (bHLH DNA-binding superfamily), and AT4G29270 (HAD superfamily).

### KEGG pathway enrichment reveals increased translation of defense response, MAPK signaling and UPR genes

In order to test how the defense response may be translationally controlled during RCNMV infection, and also to identify any unexpected translationally controlled functions, we performed KEGG analysis. In the RNA-seq dataset at 5 dpi, only the “plant-pathogen interaction” term was enriched, whereas in the Ribo-seq dataset, “plant-pathogen interaction” and “MAPK signaling pathway-plant” terms were enriched (Fig. 3A, Supplemental File S3). All the DEGs and DTGs in these pathways were upregulated. By 8 dpi, many more pathways were enriched for both RNA-seq and ribo-seq datasets, such as “Metabolic pathways”, “Plant hormone signal transduction”, and “Protein processing in endoplasmic reticulum”, among others. This demonstrates that by the time that plants become symptomatic, virus infection disrupts several cellular pathways.

**Figure 3.**
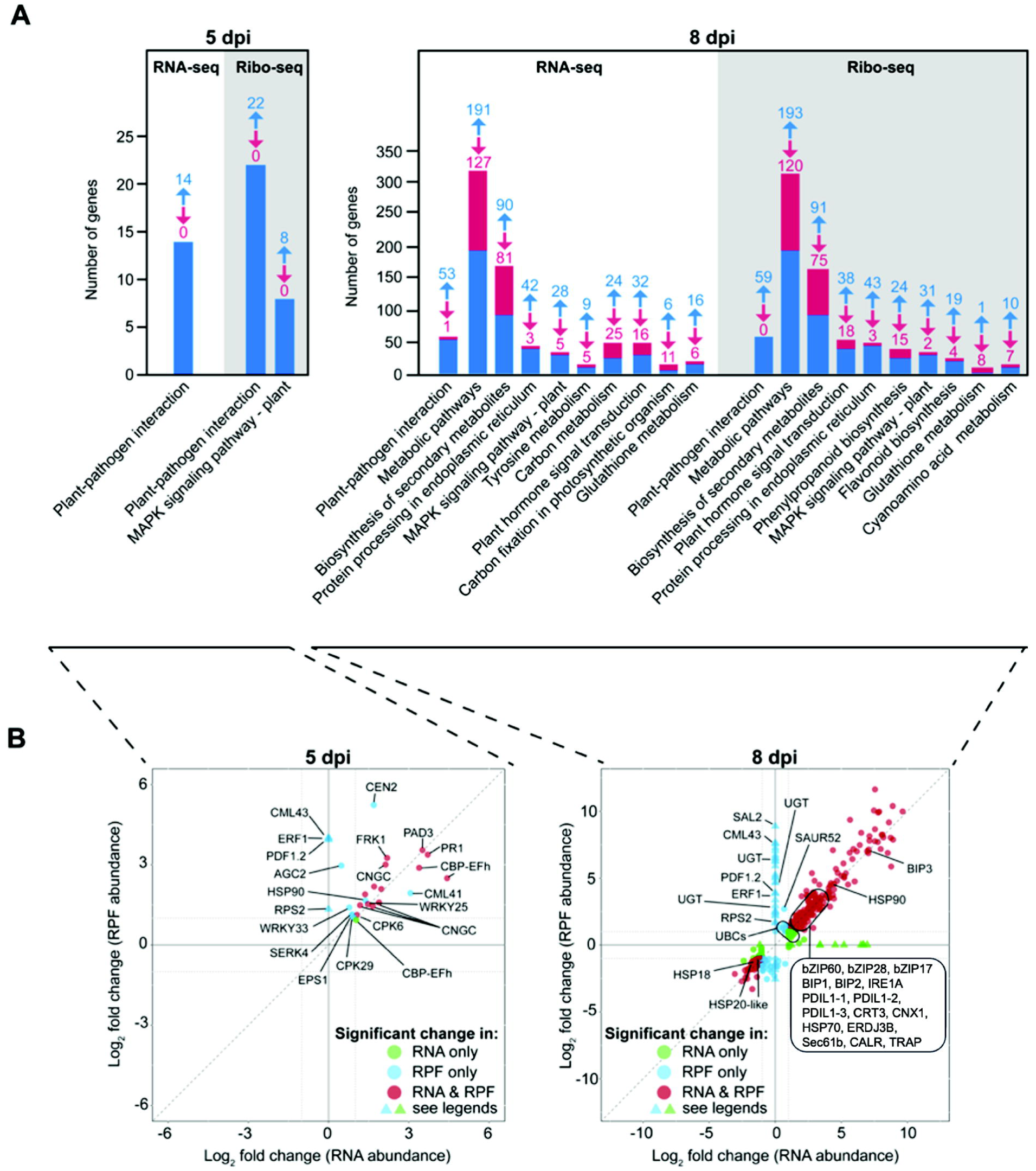
KEGG pathway enrichment analysis. **(A)** The enriched pathways were identified via KOBAS (118) using the DEGs (RNA-seq) and DTGs (Ribo-seq) at 5 and 8 dpi. The upregulated and downregulated genes were used together for the analysis. The blue and the red arrows with the number denote the number of upregulated and downregulated DEGs/DTGs, respectively, in the enriched pathway. **(B)** Scatterplot of log_2_ FC (RPF abundance) vs log_2_ FC (RNA abundance) with the DEGs/DTGs in the enriched pathway. The blue triangles refer to the DTGs that did not have a valid output (non-NA) from the RNA-seq data via DESeq2 and therefore, their log2FC (RNA abundance) was set to zero. The green triangles refer to the DEGs that did not have valid output (non-NA) from the ribo-seq data via DESeq2 and therefore, their log2FC (RPF abundance) was set to zero. (DEGs) differentially expressed genes, (DTGs) differentially translated genes, (FC) fold change.

Next, we plotted DEGs vs RPFs that were present in the enriched pathways (Fig. 3B, Supplemental File S4). At both 5 and 8 dpi, most of the genes were regulated at transcription as the RPFs changed in proportion to change in mRNA level (Fig. 3B, points near the x = y diagonal dotted line). A few other genes (denoted as blue triangles) were identified as DTGs but did not have any DESeq2 output from the RNA-seq data, mostly because of the low read counts. Because the RNA abundance did not increase detectably in the RCNMV-infected plants, but the RPF levels did increase, we speculate that these genes may be regulated at the translational level. Examples include defense-related genes such as AT3G23240 (ERF1), AT4G26090 (RPS2), AT5G44420 (PDF1.2), and AT5G44460 (CML43), which were identified only as DTGs at both the time-points.

Interestingly, the genes in the pathway “Protein processing in endoplasmic reticulum” that were identified as DEGs, with a proportional increase in RPFs, at 8 dpi were regulated only at the level of transcription. These genes included BiP1, BiP2, BiP3, PDILs, CRT, CNX, HSP70, HSP90 and ERDJ3B which are known to be upregulated downstream of IRE1-mediated bZIP60 splicing in the cytoplasm that occurs during unfolded protein response (UPR) (63, 64). The function of these canonical UPR-responsive gene products is to ameliorate the ER-stress by increasing the capacity of the ER for protein folding, import, export, and the quality control (65). As BiP and other ER-resident genes, including the UPR signalling genes bZIP60, bZIP28 and bZIP17, are upregulated at 8 dpi (Fig. 3B), we conclude that RCNMV infection elicits UPR in Arabidopsis.

### Only a small proportion of RCNMV RNAs are undergoing translation, but RCNMV is still the most translated RNA in the cell

The proportion of RCNMV-mapped reads was calculated as the number of reads that mapped uniquely to RCNMV RNAs divided by the sum of the reads that mapped uniquely to RCNMV RNAs and Arabidopsis reference genome. (See Supplemental File S1). From 5 to 8 dpi, RNA-seq reads mapping to RCNMV increased from ∼ 0.2 % to ∼ 21.5 % of total RNA-seq reads (Fig. 4A). On the other hand, the proportion of 26-29 nt ribo-seq reads mapping to the RCNMV RNAs increased from ∼ 0.02 % to ∼ 1.7 % of total ribo-seq reads (Fig. 4A). Such a high proportion of RCNMV RNA-seq reads with a relatively lower proportion of ribo-seq reads was not surprising as the RNA-seq reads includes the viral RNAs isolated from virus particles which accumulate to high levels and in which no translation occurs. At 5 dpi, only ∼0.2% of the mapped RNAseq reads were derived from RCNMV genome, but these RCNMV RNA reads were abundant relative to those mapping to any particular Arabidopsis mRNA (Fig. 4B). Unsurprisingly, RCNMV RNAs were the most abundant RNA species of any kind at 8 dpi (Fig. 4B). A similar pattern can be seen for the Ribo-seq reads (Fig. 4B). At 8 dpi, ∼1.7% of RPFs were derived from RCNMV mRNAs, indicating that the number of ribosomes translating RCNMV RNA exceeded that for any individual host mRNA (Fig. 4B). (However, the RPFs derived from chloroplast genome may be underestimated because we used only the 26-29 nt Ribo-seq reads for the analysis and the RPFs from chloroplasts are known to be longer than cytosolic RPFs (66).)

**Figure 4.**
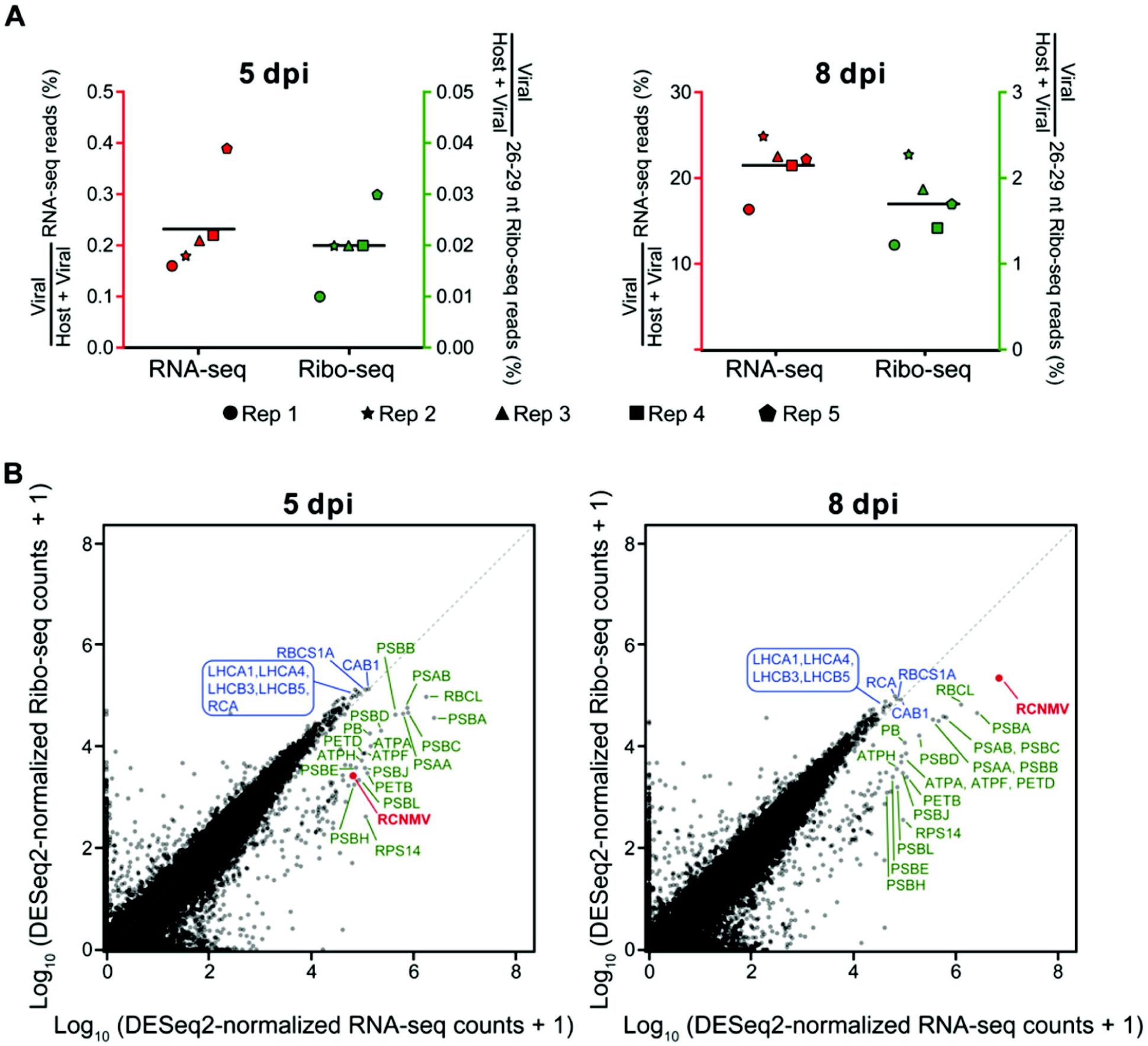
Abundance of RCNMV-mapped RNA-seq and ribo-seq reads. **(A)** Proportion of RNA-seq and ribo-seq reads in RCNMV-infected plants that map to RCNMV genome. **(B)** Scatterplot display the DESeq2-normalized read counts from RCNMV compared to that from Arabidopsis genome. Values displayed are the arithmetic means of the log10 (1+x)-transformed counts across the five biological replicates. Only the Arabidopsis genes that had higher RNA-seq read counts than RCNMV at 5 dpi are labelled in the scatterplot. RCNMV is denoted by red color. Blue-colored labels show nucleus-encoded genes, and green-colored labels show chloroplast-encoded genes.

### Negative strand but not positive strand reads are enriched in the region corresponding to sgRNA1

The integrated density of RNA-seq reads over RCNMV genome was visualized by aligning entire read sequences to the RCNMV RNA1 and RNA2 genomic RNAs from all the replicates combined (Fig. 5). Even though a previous RNA-seq experiment (54) used a different host (*Nicotiana benthamiana*), RNA extraction protocol, library preparation strategy/kit, and Illumina sequencer, we observed similar characteristics of RNA coverage profiles in this RNA-seq experiment (Fig. 5), including: (i) no greater abundance of positive sense reads corresponding to the subgenomic RNAs, CPsgRNA and SR1f, (ii) CPsgRNA negative-strand accumulation to much higher levels than negative strand reads mapping to upstream portions of RNA1, supporting the premature transcription termination model (41, 67) for generation of the sgRNA, and (iii) more positive sense reads toward the 5’ half of RNA2 than the remainder of RNA2 with an inflection point near the trans-activator (TA) region.

**Figure 5.**
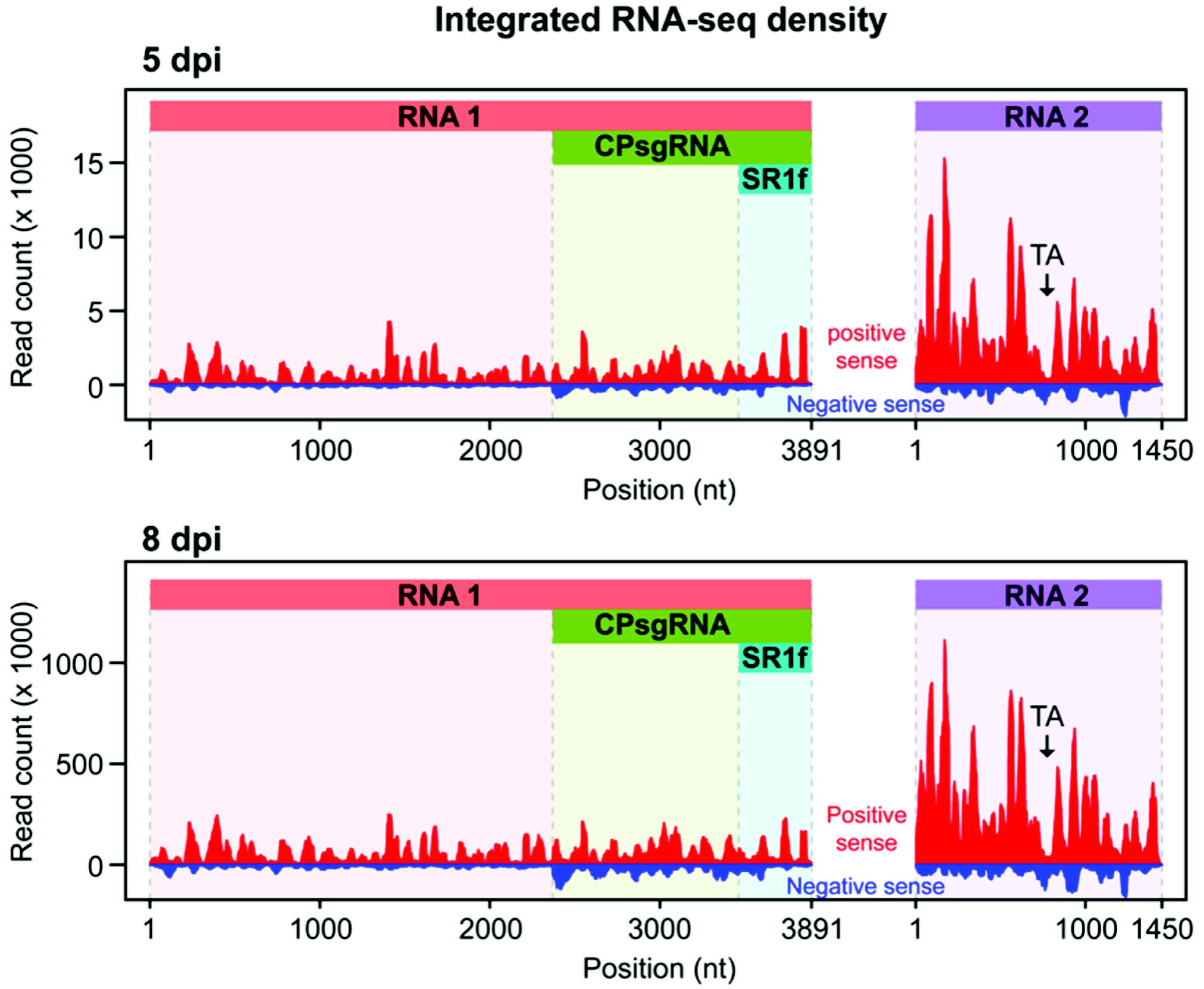
Integrated RNA-seq density over RCNMV RNA1 and RNA2. All the nucleotides of RNA-seq reads were plotted as a histogram which represents the integrated read density for all the reads combined from the five biological replicates at each time-point. The red and the blue curve denote the reads mapping to the positive-sense and negative-sense RNA, respectively. The black arrow denotes the location of the TA sequence that interacts with the TA-BS sequence of RNA1 and result in premature transcription termination of CPsgRNA. TA region coincides with the point of inflex of RNA-seq density. (nt) nucleotide, (TA) trans-activator.

### Ribo-seq profiles on RCNMV genome reveal ribosomal frameshifting rate and extremely high translation of the coat protein ORF

The 28 nt RNA-seq and Ribo-seq reads that mapped to RCNMV genome were visualized using the riboSeqR package (68). The RNA-seq and Ribo-seq profiles on RCNMV genome were reproducible across the biological replicates. Furthermore, riboSeqR correctly identified the three ORFs in RNA1 (Fig. 6, red and blue horizontal bars over each profile) and one ORF in RNA2 (Fig. 7, green horizontal bars over each profile) as being translated. Except for the increase in number of reads, we did not see any other change in coverage pattern from 5 dpi to 8 dpi in these profiles.

**Figure 6.**
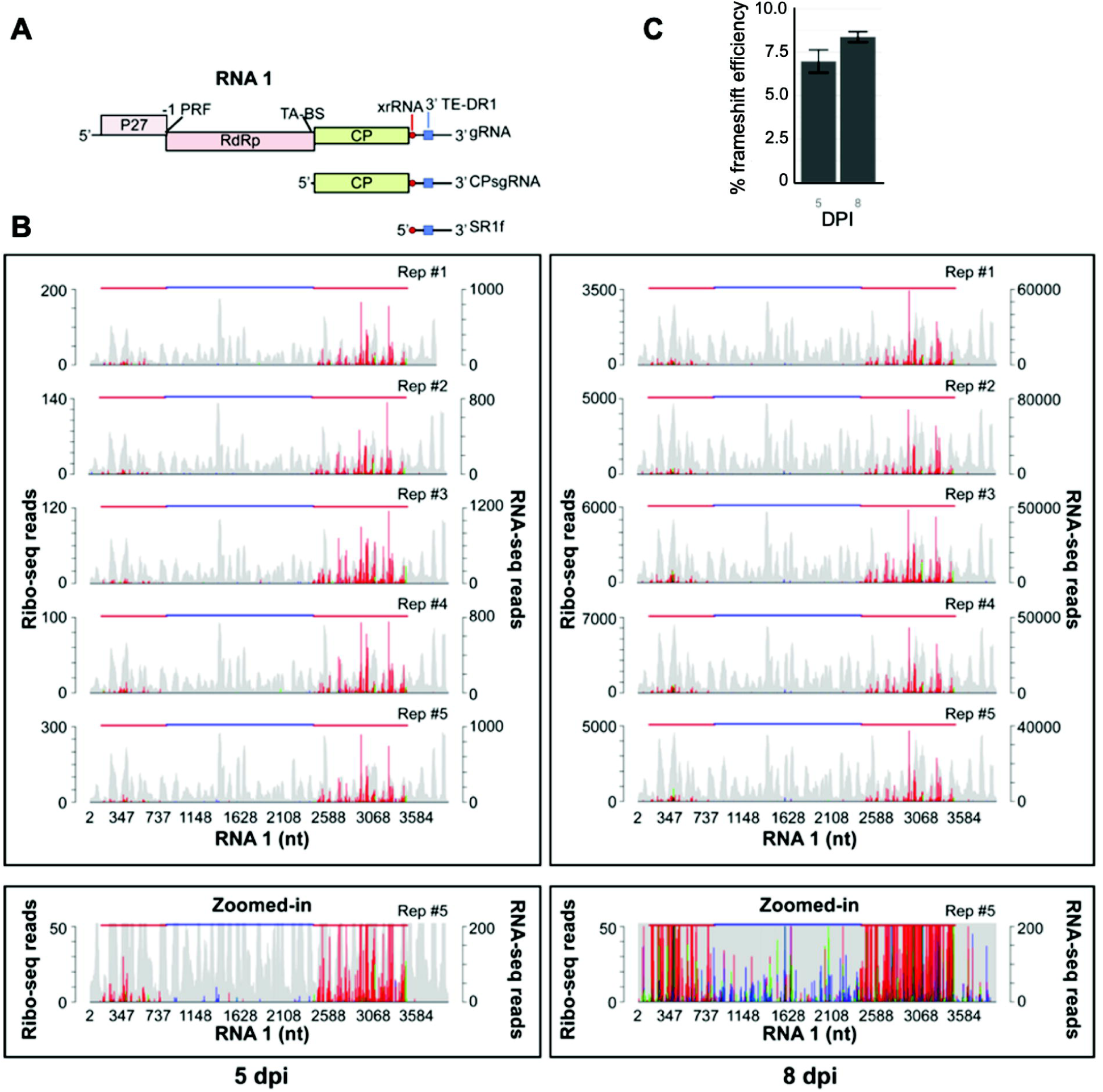
Ribo-seq and RNA-seq profiles of RCNMV RNA1. **A.** Genome map of RCNMV to guide interpretatioin of with reads mapped in panel B. **B.** RiboSeqR (53) was used for plotting the positions of the first nt of each RPF (colored) and each RNA read (gray) as a histogram. The color of the peak is determined by the translating frame (frame 0, 1, 2) to which the first nt of the RPF maps. The figure shows the profiles of only the 28-nt RPFs and RNA reads for all the five biological replicates at both time-points. The bottom panel depicts a zoomed-in image of replicate 5 to show the RPFs mapping to the RdRp ORF. The translated ORF inferred by riboSeqR (blue and red horizontal line) corresponds to the known RNA1 annotations. -1 PRF can be observed as the red frame (RPFs at p27 ORF) switches to the blue frame (RPFs at RdRp ORF) at the frameshift site. This map aligns with that in panel A. (nt) nucleotide, (RdRp) RNA-dependent RNA polymerase, (PRF) Programmed ribosomal frameshifting. **C.** Frameshift rates at 5 and 8 days post-inoculation (DPI) measured as described in text.

**Figure 7.**
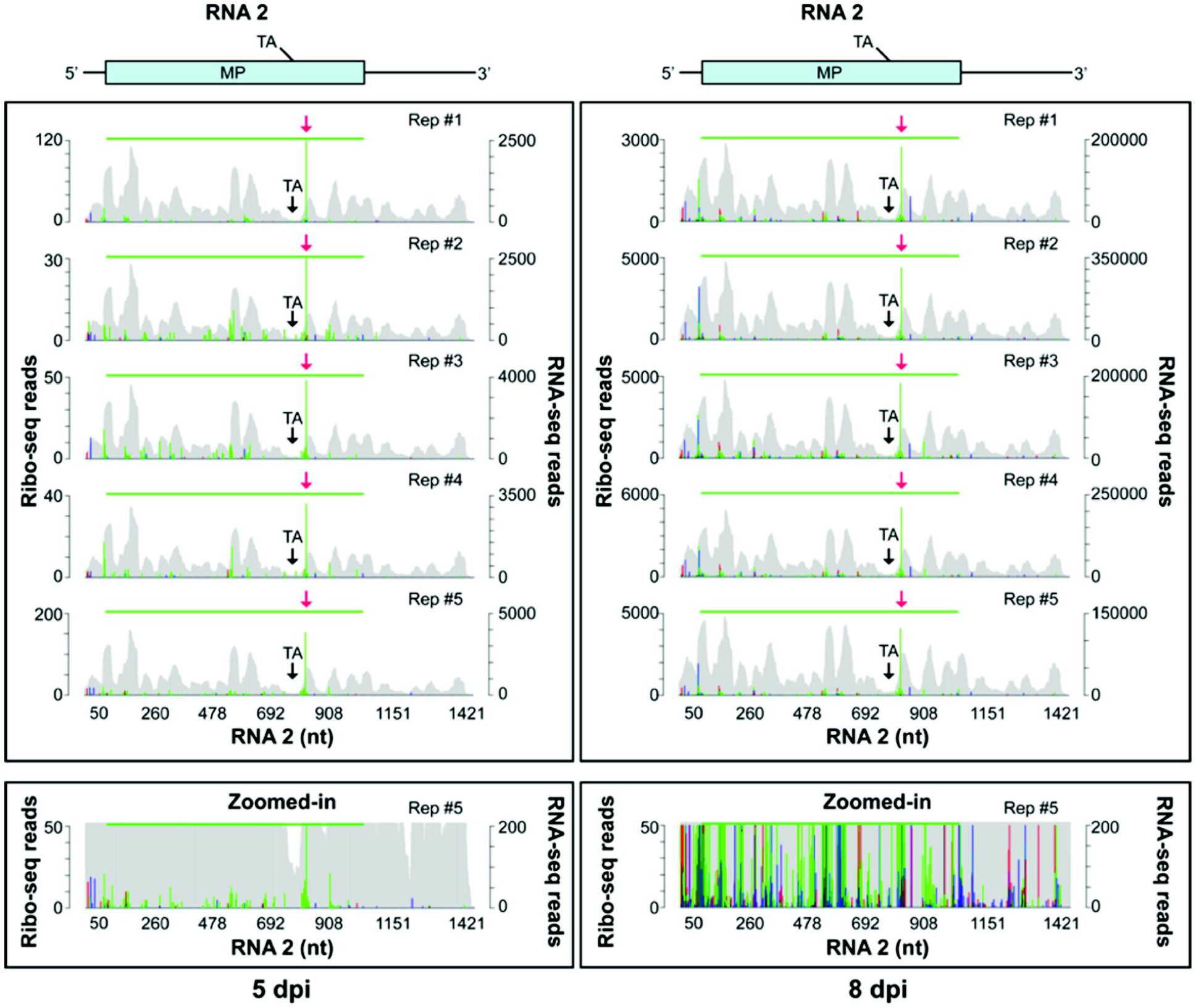
Ribo-seq and RNA-seq profiles of RCNMV RNA2. riboSeqR (53) was used for plotting the first nt of the RPFs (colored) and RNA reads (gray) as a histogram. The color of the peak is determined by the translating frame (frame 0, 1, 2) at which the first nt of the RPF maps to. The figure shows the profiles of only the 28-nt RPFs and RNA reads for all the five biological replicates at both time-points. The bottom panel depicts a zoomed-in image of replicate 5. The translated ORF inferred by riboSeqR (green horizontal line) corresponds to the known RNA2 annotation. The red arrow denotes a putative ribosomal pause site. The black arrow denotes the location of the TA sequence that interacts with the TA-BS sequence of RNA1 and results in premature transcription termination of CPsgRNA (41). TA region coincides with the point of inflex of RNA-seq density. (nt) nucleotide, (TA) trans-activator, (TA-BS) TA-binding site.

In RNA1, we detected the -1 programmed ribosomal frameshifting (-PRF) as the translating frame changes at the frameshift site, denoted by the red frame at p27 ORF and blue frame in RdRp ORF (Fig. 6A,B, see the zoomed image). By calculating the ratio of RPF per kb mapping to ORF2 to those mapping to ORF1, we calculated a frameshift rate of ∼7.5% at 5 dpi and ∼8.0% at 8 dpi (Fig. 6C). Whether this difference between timepoints is biologically relevant is unclear, but to our knowledge this represents the first direct calculation of ribosomal frameshift efficiency on a replicating plant viral RNA.

The abundance of positive sense RNA-seq reads across RNA1 is fairly uniform, suggesting low accumulation of CPsgRNA as a proportion of total viral RNA (Fig. 6B). However, the number of ribo-seq reads mapping to CP ORF is vastly higher than to the ORFs 1 and 2, indicating the CP is translated with very high efficiency (Fig. 6). In RNA2, RPF read density was very low showing that RNA2 is not translated efficiently (Fig. 7). Interestingly, we observe an extremely high RPF peak (Fig. 7, red arrow) in the MP ORF that was present in all the samples at both time-points. This may reveal a strong ribosomal pause site.

## Discussion

### Translational response to RCNMV infection

Unlike many vertebrate viruses, plant viruses do not induce a global shutdown of translation. However, it is clear that many mRNAs are specifically altered in translational efficiency in response to plant virus infection, including by RCNMV. A level of inhibition of host translation may arise due to viral RNAs outcompeting host mRNAs for the ribosomes, via the highly efficient cap-independent translation element (BTE) present in the 3’ UTR of RNA1 (45). The BTE binds the rate-limiting translation initiation factor, eIF4F with high affinity (47). Its presence in RCNMV genomic RNA, CPsgRNA1 and the noncoding SR1f sgRNA may allow these RNAs to sequester enough eIF4G to cause a reduction in host mRNA translation (14). However, many if not most sampled cells may not have had sufficient available BTE to affect host translation. Instead, the translational control we see may be due to plant defense signalling. We sampled non-inoculated, systemically infected tissue, and owing to the uneven, patchy nature of virus infection, it’s likely that the sampled tissue contained numerous uninfected cells, and cells either too early in infection to have accumulated enough viral RNA to affect host translation (especially at 5 dpi) or so late in infection that all the viral RNA is packaged in virions (especially at 8 dpi) and inaccessible to translation machinery. Thus, the changes in translation efficiency we detected may be host defense signalling responses, including in uninfected cells that have received the signal prior to virus entry. We must also bear in mind that the defense response is reduced by the dcl2/dcl4 knockout, which eliminates antiviral RNA silencing and can have effects on the NLR and salicylic acid-mediated defense pathways, but do not prevent them from functioning (69–71).

It appears that the early translationally-regulated response to virus systemic infection is specific and targeted towards ameliorating the virus infection. It makes sense that we see translational induction of immunity genes at the very earliest stages of infection, and probably in cells not yet infected, as the translational response is much more rapid than transcription. In contrast, in the 8 dpi tissue, the massive increase in transcription of many mRNAs presumably allowed much higher expression of defense-related genes, and clearly involves many additional gene functions. Many of these changes in gene expression later in infection may be by-products of dysregulated pathways resulting from the plant being very sick. Given that the infected plants lacked functional DCL2/4 genes, it is noteworthy that defense genes were induced in the absence of the RNA interference response. This reveals the redundancy of plant defense responses to virus infection.

One mRNA (AT4G09770) that remained translationally but not transcriptionally upregulated at both timepoints encodes a Tumor necrosis factor receptor-associated factor (TRAF)-like protein (Fig. 2B). In mammals, there are seven TRAF proteins that interact with various cell surface receptors via the TRAF-domain; they play key roles in the immune response and apoptosis (72). Arabidopsis genome has more than seventy TRAF domain-containing proteins (73). AT4G09770 belongs to the TRAF/MATH-only class of TRAF proteins that are involved in regulating autophagy, plant immunity, and gametophyte development (73). Two other TRAF-proteins belonging to the same class, MUSE13 and MUSE14, form a TRAFasome that degrades NLR immune receptors, SNC1 and RPS2, via modulating ubiquitination (74). During RCNMV infection, whether AT4G09770 (TRAF-like family protein) is translationally-upregulation by the plant to prevent overaccumulation of plant NLR proteins or by the RCNMV as a counter-defense strategy remains to be determined.

By 8 dpi, multiple genes in the UPR pathway were transcriptionially induced at high levels. The elicitation of UPR pathways has been shown in a variety of plant-virus infections (65, 75). Since RCNMV forms virus replication complex at the ER by restructuring the ER membrane (76, 77), it is very likely that it upregulates the genes to increase the ER membrane surface area. Over-accumulation of viral proteins can also elicit UPR that serves to alleviate ER-stress related cell death (75). Similar to our previous report on unfolded protein response in maize seedlings (78), we observed that during RCNMV infection, the UPR genes are upregulated only transcriptionally (Fig. 3B). The upregulation of UPR genes support the findings in a previous report that showed an increased amount of BiP proteins that are associated with the ER during RCNMV infection (76). It still needs to be investigated whether UPR elicitation during RCNMV infection acts as a (i) proviral mechanism to increase virus replication by increased formation of virus replication complexes at the ER, (ii) proviral mechanism to increase the protein folding capacity of the ER for efficient synthesis of functional viral proteins, (iii) proviral or plant’s prosurvival pathway to reduce ER stress-mediated cell death, or (iv) antiviral defense response to limit RCNMV infection.

### Viral translation

Mapping of RPFs to viral ORFs revealed, as expected, the change in reading frame (red to blue, Fig. 6) at the ribosomal frameshift site, and large reduction in RPFs mapping to ORF2 relative to ORF1 due to the infrequency of ribosomal frameshift. Also, the coat protein is translated with far more efficiency than the nonstructural proteins, which is expected as this virus requires 180 CP subunits per virion which contains one genomic RNA. As one RdRp can likely replicate multiple RNAs, less than one RdRp molecule would be needed per viral RNA, hence orders of magnitude less RdRp than CP is required for a productive infection. These extreme ratios are reflected in the RPF counts for the RdRp and CP-encoding ORFs.

Unexpectedly, here (Fig. 5) and in a previous study (54) we did not detect significantly more RNAseq reads mapping to the 3’ end of the genome that encodes the CP. We expected more reads, due to accumulation of CPsgRNA1 (41). However, inspection of previous northern blot hybridizations revealed very low levels of CPsgRNA1 relative to genomic RNA (41, 54). Thus, given the huge ratio of CP ORF RPFs per RNAseq read, the CP ORF must be translated with extremely high efficiency (Fig. 6B). To facilitate this high level of translation, it is possible that the CP ORF is also translated directly from genomic RNA. This would require an internal ribosome entry site (IRES) upstream of the CP AUG. Indeed, internal initiation of translation of the CP ORF has been detected in other tombusvirids such as turnip crinkle virus (79) and MCMV (80), indicating some IRES activity in the noncoding region upstream of the CP ORF in those viruses. In contrast, any IRES activity in RCNMV would have to be located in the coding region of ORF2, because ORF2 terminates just one nucleotide upstream of the ORF3 (CP ORF) start codon. This region at the 3’ end of ORF2 includes the *cis*-acting sequence that base pairs with the *trans*-activator (TA) RNA sequence in RCNMV RNA2 to facilitate production of CPsgRNA1 via premature termination of the RdRp (41). Thus, if an IRES is present, the 3’ end of ORF2 would perform three functions: (i) coding for the C terminal portion of the RdRp, (ii) providing the RNA structure and sequence for interaction with the TA of RNA2, and (iii) serving as an IRES. Another, explanation is that, if the vast majority of RNA1 is packaged, the ratio of CPsgRNA1, which is not encapsidated, to unencapsidated RNA1 may be much higher than to the total measured RNA1 (encapsidated + unencapsidated). Both of the above explanations may apply simultaneously.

Sequence and structural requirements for frameshifting on plant viral RNAs has been studied in depth using either cell-free translation of the viral genomic RNA, or in plant cells (protoplasts) which requires use of reporter genes or immunoblotting to detect the translation products (81–83). While valuable for observing effects of mutations on the relative level of frameshifting, these methods, which reflect steady state protein levels, are less direct than ribo-seq, which reveals the quantities of ribosomes on the pre- and post-shift ORFs and thus the actual rate of frameshifting on replicating viral RNA in an infected cell. However, the frameshift rate of 7.5 to 8 percent that we calculated by measuring ribosome occupancy of each ORF (Fig. 6C) agrees with that calculated by Kim and Lommel (7.4 ± 2.9%) in uninfected protoplasts using a non-replicating single reporter construct lacking a second reporter as an internal control (84). Thus, the ability to replicate and the changes to the cellular environment due to virus infection may not affect RCNMV frameshift efficiency, as also concluded by Tajima et al. (83).

Previously, it was considered that ribosomal pausing at the frameshift site would be required for ribosomal frameshifting (85, 86), although, more recently, ribo-seq on retroviruses (87), coronaviruses (22, 88) and encephalomyocarditis virus (EMCV) (89) failed to detect such a pause at the -1 frameshift site. Similarly, we did not detect any ribosomal pausing (localized spike of a high number of RPFs) at the frameshift site (Fig. 6B). This is consistent with observations using different methods that showed no correlation between ribosomal pausing and frameshifting (90). In contrast, a modified nuclease digestion (disome footprint profiling) with SARS-CoV-2-infected cells did show ribosome pausing at the frameshift site and this observation was verified using cryo-EM (91). Therefore, whether ribo-seq is suitable for assessing ribosomal pausing during frameshifting is still debatable.

Finally, an unexpected observation was the massive accumulation of reads at one site in the MP ORF on RNA2 (Fig. 7). This may reflect a strong pause to allow proper folding of the nascent MP strand before synthesis of the entire protein is complete. Pausing occurs often during protein synthesis, especially on mRNAs encoding membrane proteins to allow entry into the ER (92). Presence of consecutive proline codons is known to induce ribosomal pausing owing to the slower time for peptide bond formation by prolines (93, 94), and the kink induced in the polypeptide chain by prolines can slow the movement of the nascent peptide through the exit tunnel of the ribosome (95). Proline codon doublets occur at nts 757-762, 796-801, and 871-876 in the MP ORF. A positively charged (RK-rich) tract of amino acids preceding two or more consecutive proline residues is particularly effective in stalling the ribosome, as highly charged tracts of nascent polypeptide move slowly through the ribosome exit tunnel (96). Such a situation exists in the RCNMV MP ORF at the site of the large peak of RPFs. Bases 820-840 encode RKVPKRK followed 11 codons downstream by the PP doublet at nts 871-876. Given that the exit tunnel accommodates about 30 amino acids of nascent chain, the highly basic tract (which also includes a proline) would be in the exit tunnel as the PP peptide bond formation takes place in the A and P sites.

It is noteworthy that very few RNAseq reads map to the TA stem-loop of RNA2 (nts 779-811, with nts 790-797 in the loop that base pairs to RNA1) in our RNA coverage profile (Figs. 5 and 6). This is just upstream of the potential ribosomal pause site. We speculate that ribosomal pausing may lead to stacking of stalled ribosomes that can elicit No-Go decay, resulting in an endonucleolytic cleavage upstream of the pause site (97, 98). If this hypothesis is true, it would explain the unusual RNA2 RNAseq coverage pattern, and low translation of RNA2 as evident by sparsely distributed RPFs over RNA2 (Fig. 7). However, the known functions of the TA, as a cis-acting RNA2 replication element (99), and to generate CPsgRNA1 by base-pairing to RNA1 (41) would take place in replication vesicles thought to be sequestered from ribosomes, to prevent ribosomes from competing with the replicase for the viral RNA template (52, 100). Thus, the ribosome pausing event that generates the unusual RPF pattern may be physically separate from these TA functions. Finally, Mizumoto *et al.* (101) have shown that translation of RNA2 depends on replication, suggesting RNA modification during replication or co-colocalization of replication and translation components are required. This may affect the distribution abundance and distribution of RPFs in ways that aren’t obvious.

In conclusion, ribosome profiling has allowed us to identify unexpected translational response of host genes, accurate measurement of viral translation efficiency, while the exact effects of the noncoding SR1f on translation require future experiments, perhaps with more synchronous infection systems.

## Materials and Methods

### *In vitro* transcription of RCNMV RNAs

RCNMV plasmid constructs used for *in vitro* transcription were described previously (102). pRC169c and pRC2|G are cDNA clones with T7-promoter for *in vitro* transcription of infectious RCNMV RNA1 and RNA2, respectively. One µg of SmaI-linearized pRC169c and pRC2|G were used as template for *in vitro* transcription using MEGAscript T7 Transcription kit (Invitrogen AM1334) followed by DNase treatment according to manufacturer’s protocol. The transcription reaction was carried out at 37°C for 4 h and DNase treatment at 37°C for 30 min. Subsequently, RNA was purified using Zymo RNA clean & concentrator -5 kit (Zymo Res. R1015) and eluted in nuclease-free water.

### RCNMV-infected sap preparation

*N. benthamiana* plants were grown in a growth chamber with 16 h light at 24°C and 8 h dark at 20°C. Four-week-old plants were mechanically inoculated with (i) 10 mM sodium phosphate (pH 6.8) buffer (Mock-inoculated), or (ii) 1 µg RNA1 plus 1 µg RNA2 in 10 mM sodium phosphate (pH 6.8) buffer per leaf (RCNMV-inoculated) and the growth conditions were changed to 12 h light at 20°C and 12 h dark at 20°C. After the appearance of symptoms, RT-PCR was used to verify RCNMV replication in the non-inoculated systemic leaves. Subsequently, the leaves were ground in 10 mM sodium phosphate (pH 6.8) buffer using mortar and pestle and the resulting sap was used for inoculating Arabidopsis plants.

### Inoculating Arabidopsis with RCNMV

Arabidopsis double knock-out mutant line, *dcl2-1/dcl4-2t* (Germplasm CS66078) (Xie et al. 2005) was obtained from the Arabidopsis Biological Resource center (abrc.osu.edu) and the T-DNA insertion was verified by genotyping (O’Malley et al. 2015). The plants were grown in growth chambers with 16 h light at 24°C and 8 h dark at 20°C. After 3 weeks, the growth condition was changed to 12 h light at 20°C and 12 h dark at 20°C. Symptomatic leaves from RCNMV-inoculated *N. benthamiana* plants or healthy leaves from mock-inoculated *N. benthamiana* plants were ground with a mortar and pestle in 10 mM sodium phosphate (pH 6.8) buffer, and the resulting sap extract was rubbed on 2-3 large rosette leaves per Arabidopsis plant (4-week-old) using Q-tips and carborundum.

### RT-PCR and qRT-PCR

For the time-course experiment, we harvested 3-4 of the youngest (non-inoculated) leaves per plant at growth stages 3.7 and 5.1 as described by Boyes et al. (103). These leaves from 3 plants were pooled to make 1 biological replicate and 3 biological replicates were collected at 4, 5, 6, 7, 8, 9, and 10 dpi for RCNMV-infected plants and at 10 dpi for mock-infected plants. The leaves were pulverized in a tissue lyzer at 1500 rpm for 30 s (twice). Total RNA was extracted using Zymo Direct-zol RNA Miniprep kit (Zymo Res. R2050) according to the manufacturer’s protocol followed by quantification using Qubit RNA HS assay kit (Invitrogen Q32852). Total RNA integrity was verified by agarose gel electrophoresis. RevertAid First Strand cDNA Synthesis kit (Thermo Scientific K1622) was used for DNase treatment and cDNA synthesis (with random hexamers) according to the manufacturer’s protocol.

For RT-PCR, a 10 µl PCR reaction mix was prepared with 5 µl GoTaq G2 green master mix (Promega M7823), 1 µl 10-fold diluted cDNA template, 200 nM each of RCNMV_1457_FP (5’-CAACAGGGCTCAAGGGAGAG-3’) and RCNMV_1574_RP primers (5’- GAATTTGAGGGCATCGACGC-3’). The PCR conditions were as follows: 98°C (2 min); 30 cycles of 98°C (10 s), 60°C (15 s), 72°C (15 s); 72°C (2 min); 4°C hold.

For qRT-PCR, a 10 µl qPCR reaction was prepared with 1x iQ SYBR Green Supermix (Bio Rad 1708880), 300 nM each of forward and reverse primers, and 1 µl 10-fold diluted cDNA template. The qPCR runs were carried out in 384-well plates with 3 technical replicates per sample in a Bio Rad CFX384 system with the following reaction conditions: 95°C for 3 min (Polymerase activation and DNA denaturation), 40 cycles of 95°C for 10 s (Denaturation), 60°C for 60 s (Annealing, extension/Plate reading) followed by melt curve analysis (55-95°C, 0.5°C increment, 5 s). AtSAND and AtPDF2 genes were used as reference genes with the primer pairs Athal_PDF2_FP (5’-TCATTCCGATAGTCGACCAAG-3’) plus Athal_PDF2_RP (5’-TTGATTTGCGAAATACCGAAC-3’) and Athal_SAND_FP (5’-GTTGGGTCACACCAGATTTTG-3’) plus Athal_SAND_RP (5’-GCTCCTTGCAAGAACACTTCA-3’). These genes showed unchanged expression in response to infection by a variety of viruses (Lilly et al. 2011). Prior to using these as reference genes, we verified their consistent expression between our experimental conditions. The primer efficiency calculation, ΔΔCt calculation, and statistical analysis were performed using the Bio Rad CFX manager software.

All the primers were synthesized by Integrated DNA Technologies (IDT) and purified by standard desalting.

### Ribosome profiling

Ribosome profiling protocols from (57, 78, 104, 105) were used with modifications as follows.

#### Lysate preparation

For ribosome profiling, 5 biological replicates were collected each for mock- and RCNMV-infected plants at 5 and 8 dpi. Each biological replicate was comprised of 3-4 young leaves per plant pooled together from 18 plants. Consistent with the time-course experiment, the symptoms appeared after 7 dpi (Fig. 1A). RCNMV infection was verified by RT-PCR (Fig. 1B). The tissues were collected in 50 ml falcon tubes and coarsely pulverized by vigorously shaking the tube with two 4.8 mm metal beads in it. Aliquots of coarsely ground tissues were finely pulverized in 1.5 ml tubes using tissue lyzer at 1500 rpm for 30 s (twice) followed by the addition of 800 µl of polysome extraction buffer (PEB; 20 mM Tris-HCl pH 7.5, 140 mM KCl, 5 mM MgCl_2_, 1% (v/v) Triton X-100, 0.5% (v/v) Igepal CA 630, 146.1 mM sucrose, 100 µg/ml cycloheximide, 100 µg/ml chloramphenicol, 0.5 mM DTT, 0.5 µl/ml Turbo DNase (Invitrogen AM2238), and EDTA-free protease inhibitor (Thermo Scientific A32965)). The crude lysate was incubated in ice for 20 min on a rocker followed by clarification using two rounds of centrifugation (21,100 x g, 15 min, 4°C). Subsequently, the absorbance of the lysate was adjusted with PEB to A_254_ (Lysate - PEB) ∼ A_260_ (Lysate - PEB) ∼ 6. 400 µl of the absorbance-adjusted lysate was used for ribosome profiling and 200 µl for RNA sequencing.

#### RNase1 digestion

400 µl lysate was centrifuged (21,100 x g, 5 min, 4°C) to remove any remaining debris. Subsequently, 2 µl RNAse1 (Invitrogen AM2295) was added and the RNase digestion was carried out for 60 min at 28°C in a thermomixer at 400 rpm. The tubes were immediately transferred to ice and 5 µl Superase-IN (Invitrogen AM2696) was added to terminate the RNase digestion reaction.

#### Pelleting the monosomes

To the RNase-treated lysate, PEB was added to make the total volume ∼750 µl and was carefully layered on a precooled 350 µl sucrose cushion (35% (w/v) sucrose, 20 mM Tris-HCl pH 7.5, 140 mM KCl, 5 mM MgCl_2_, 100 µg/ml cycloheximide, 100 µg/ml chloramphenicol, 0.5 mM DTT, 5 µl/ml Superase-IN (Invitrogen AM2696)) in mini-ultracentrifuge tubes (Thermo Scientific 45237) followed by ultracentrifugation at 57,000 rpm (131,500 x g) for 90 min at 4°C with slow acceleration and deceleration in a Sorvall mini ultracentrifuge (Discovery M150) with S150-AT fixed angle rotor (Thermo Scientific 45582). Subsequently, the supernatant was removed and the pellet was carefully rinsed with 500 µl nuclease-free water.

#### Ribo-seq RNA purification

600 µl proteinase-K buffer (10 mM Tris-HCl pH 7.5, 1% SDS, 200 µg/ml proteinase K (Thermo Scientific EO0491)), prewarmed to 42°C, was added to the monosome pellet and incubated at room temperature (RT) for 5 min, resuspended the pellet by pipetting, transferred to 1.5 ml microfuge tubes, and incubated at 42°C for 30 min. Subsequently, it was heated to 65°C for 2 min and immediately proceeded to RNA purification using hot acid-phenol chloroform method as follows. 600 µl acid-phenol chloroform (5:1, pH 4.5, Invitrogen AM9720), prewarmed to 65°C, was added and mixed, incubated at 65°C for 5 min, mixed intermittently, incubated in ice for 5 min, centrifuged (21,100 x g, 2 min), transferred aqueous phase to new tubes, added 600 µl RT acid-phenol chloroform, mixed, incubated at RT for 5 min, centrifuged, and transferred ∼500 µl aqueous phase to new tubes. Subsequently, 500 µl chloroform-isoamyl alcohol (24:1, Sigma-Aldrich 25666) was added and mixed, incubated at RT for 1 min, centrifuged and ∼400 µl aqueous phase was transferred to a 1.5-ml tube containing 2 µl Glyco Blue (Invitrogen AM9516) and 45 µl 3M sodium acetate pH 5.5 (Invitrogen AM9740). RNA was precipitated by the addition of ∼450 µl ice-cold 100% isopropanol and overnight incubation at -80°C. The RNA pellet was collected by centrifugation (21,100 x g, 45 min, 4°C), washed twice with 1 ml ice-cold 80% ethanol, air-dried and resuspended in nuclease-free water followed by Nanodrop quantification.

#### DNase treatment

A 50 µl reaction with 10 µg RNA, 1 µl Turbo DNase (Invitrogen AM2238), and 5 µl 10x Turbo DNase buffer was incubated at 37°C for 30 min, followed by addition of 1 µl more Turbo DNase and incubation at 37°C for another 30 min. DNase-treated RNA was purified using Zymo RNA clean & concentrator -5 kit (Zymo Res. R1015) according to the manufacturer’s protocol, except with one modification (1.5 volume of ethanol was used instead of 1 volume). Clean DNase-treated RNA was eluted in nuclease-free water and quantified using Nanodrop. Quality of RPFs was assessed by electrophoresis of denatured RNA in a 15% TBE-Urea gel (Invitrogen EC6885BOX) at 120 V for 5 min and 200 V for 75 min, stained the gel with SYBR gold (Invitrogen S11494) and determined the sharpness of the RPF band between the 28 nt *(5’-AUGUACACGGAGUCGACCCGCAACGCGA-3’, Sigma, HPLC purified)* and the 34 nt *(5’-AUGUACACGGAGUCGAGCUCAACCCGCAACGCGA-3’, Sigma, HPLC purified)* RNA oligos.

#### rRNA-depletion

A half-reaction of Ribo-Zero for plant seed/root kit (Illumina MRZSR116) was used per ∼5 µg of DNAse-treated RNA according to the manufacturer’s protocol, except with one modification (the 50°C incubation step was not performed for Ribo-seq samples). Subsequently, rRNA-depleted RNA was purified using Zymo RNA clean & concentrator -5 kit (Zymo Res. R1015) according to modified Zymo protocol described above and eluted in 11 µl nuclease-free water.

#### Size-selection

Denatured rRNA-depleted RNA was subjected to electrophoresis on a 15% TBE-Urea gel (Invitrogen EC6885BOX) at 120 V for 5 min and 200 V for 75 min, stained with SYBR gold (Invitrogen S11494) for 1 min, and visualized on a blue light transilluminator. A mix of 28 nt and 34 nt RNA oligos was run as size markers. The gel slice corresponding to the region between the bottom of 28 nt and bottom of 34 nt RNA size markers was excised and transferred to a 0.5 ml tube with a hole at the bottom made by an 18 G syringe needle, and the tube was placed in a 2 mL microfuge tube. The tube was then centrifuged (21,100 x g, 2 min, 4°C) to crush the gel slice and transfer its contents to the 2 mL microfuge tube. 500 µL RNA gel extraction buffer (0.3 M NaOAc pH5.5, 1 mM EDTA pH 8, 10 mM Tris-HCl pH 7.5, 0.25% (w/v) SDS) was added and incubated overnight at 4°C on a shaker. Eluted RPFs were filtered through 0.22 µ SpinX cellulose acetate filter columns (Sigma-Aldrich CL8161). 2 µL Glyco Blue (Invitrogen AM9516) and equal volume of ice cold 100% isopropanol were added to the supernatant and RPFs were precipitated overnight at -80°C. The RNA pellet was collected by centrifugation (21,100 x g, 45 min, 4°C), washed twice with 1 ml ice-cold 80% ethanol, air-dried and resuspended in 3.25 µl nuclease-free water.

#### Library preparation

Prior to library preparation, RPF ends were repaired using T4PNK kit (Thermo Scientific EK0031) as follows. 3.25 µl RNA was incubated at 70° C for 3 min, transferred to ice, followed by the addition of 0.5 µl 10x T4 PNK buffer A (no ATP), 0.25 µl Superase-IN, 0.5 µl T4PNK enzyme. The reaction was incubated at 37°C for 30 min, 0.5 µl 10 mM ATP added, incubated at 37°C for 1 h, then 5.5 µl nuclease-free water added, and deactivated by incubation at 75°C for 10 min and transferred to ice.

Subsequently, cDNA libraries were prepared using NEXTflex Small RNA-Seq kit v3 (Perkin Elmer NOVA-5132-05) according to manufacturer’s protocol with the following specifications: The 3’ and 5’ 4N adapters were diluted 1/3-fold for step A and step D, incubation in step A was carried out overnight, bead clean-up for step F was conducted according to the “No size-selection” protocol (https://perkinelmer-appliedgenomics.com/nextflex_small_rna_v3_no_size_selection_supplement-2/). 11 cycles of PCR were performed with barcoded primers and the final libraries were cleaned using the PAGE size selection and clean-up (Step H2) according to the manufacturer’s protocol. The quality of the libraries was assessed using an Agilent bioanalyzer high sensitivity DNA Assay kit. Libraries were quantified using Qubit dsDNA HS Assay kit (Invitrogen Q32854), diluted, pooled together, and sequenced at the Iowa State University DNA Sequencing Facility using the Illumina NovaSeq 6000 on the two lanes of S1 flow cell to yield 100-bp single-end reads.

### RNA Sequencing

200 µl lysate from the five biological replicates prepared above was used for total RNA extraction. Most of the steps are same as that for Ribo-seq with few modifications listed below.

#### Total RNA purification

Prewarmed 400 µl proteinase-K buffer was added to 200 µl lysate and incubated at 42°C for 30 min. Subsequently, it was heated to 65°C for 5 min and immediately proceeded to RNA purification using hot acid-phenol chloroform method as described for Ribo-seq, except three rounds of extraction was done using the acid-phenol chloroform, instead of two (two rounds with 65°C incubation and one round with room temperature incubation).

#### DNase treatment

RNAs were DNase treated the same as the Ribo-seq samples. DNase-treated RNA was purified using Zymo RNA clean & concentrator -5 kit (Zymo Res. R1015) according to the manufacturer’s protocol without any modifications. Total RNA integrity was verified by electrophoresis of denatured RNA in a 15% TBE-Urea gel at 120 V for 5 min and 200 V for 50 min.

#### rRNA-depletion

Half-reaction of Ribo-Zero for plant seed/root kit (Illumina MRZSR116) was used per ∼5 µg of DNAse-treated RNA according to the manufacturer’s protocol without any modifications. Subsequently, rRNA-depleted RNA was purified using Zymo RNA clean & concentrator -5 kit (Zymo Res. R1015) according to manufacturer’s protocol and eluted in 11 µl nuclease-free water.

#### Alkaline fragmentation

10 µl 2x fragmentation buffer (2 mM EDTA pH 8, 12 mM Na_2_CO_3_, 88 mM NaHCO_3_) was added to 10 µl rRNA-depleted RNA and incubated at 95°C for 20 min followed by immediate addition of 280 µl stop solution (0.3 M NaOAc pH5.5, 53.6 µg/mL Glyco Blue (Invitrogen AM9516)) and 750 µl ice-cold 100 % ethanol. After overnight precipitation at -80°C, the pellet was collected and washed as described above.

#### Size-selection

Denatured rRNA-depleted RNA was subjected to electrophoresis on a 15% TBE-Urea gel at 120 V for 5 min and 200 V for 50 min, stained with SYBR gold for 1 min, and visualized on a blue light transilluminator. 50 ng of single-stranded DNA ladder (IDT 51-05-15-01) was run as size markers. The gel slice corresponding to the region between the top of the 30 nt band and top of the 40 nt band of the IDT ladder was excised and transferred to a 0.5 ml tube with a hole at the bottom made by an 18 G syringe needle, and the tube was placed in a 2 ml microfuge tube. RNA was eluted from the gel slice using the same protocol as described for Ribo-seq samples.

#### Library preparation

See description for for Ribo-seq samples. RNA-seq libraries were pooled and sequenced using the Illumina NovaSeq 6000 on the two lanes of SP flow cell to yield 100 bp single-end reads.

### Ribo-seq and RNA-seq data analysis

#### Data preprocessing

The sequencing data from both lanes were concatenated for each sample. The quality of raw sequencing reads was assessed using FastQC v.0.11.7 (Andrews 2010). The adapters were removed from raw sequencing reads (parameters: “-a TGGAATTCTCGGGTGCCAAGG --discard-untrimmed --minimum-length 23”), further processed to trim the four random bases (parameters: “-u 4 -u -4”), that were added to both ends of Ribo-seq and RNA-seq reads during library preparation, using Cutadapt v.2.5 (106). The raw sequence data was bioinformatically depleted of rRNA, tRNA, and snoRNAs, and subsequently aligned to Arabidopsis TAIR 10 reference genome and RCNMV genome. The read statistics, such as the number and proportion of raw, processed, and aligned reads can be found in Supplemental file S1. The read length distribution of Arabidopsis genome-mapped and RCNMV positive-strand-mapped ribo-seq reads peaked at 28 nt, unlike RCNMV negative strand-mapped ribo-seq reads (Supplemental Fig. S2A). In contrast, the RNA-seq read-lengths had a much wider size distribution (Supplemental Fig. S2A). RiboToolkit was used to assess the triplet periodicity for reads of different lengths which showed that 28-nt ribo-seq reads displayed the most extreme triplet periodicity followed by 29-nt, 27-nt, and 26-nt reads (Supplemental Fig. S2B).

#### Quality control plots

The quality characteristics of ribo-seq and RNA-seq data were assessed using RiboToolkit (107), RiboTaper (108) and custom scripts. RiboToolkit (107)was used (parameters: RPF length 26-29 nt, Mismatch allowed 2, Max multi-map 1) to determine the RPF-lengths with good triplet periodicity (Supplemental Fig. S2; Mock rep #1 sample as a representative result), assess the frame-enrichment (Supplemental Fig. S2; all the samples), and the distribution of reads (Supplemental Fig. S2B; all the samples) over different mRNA features. The data was obtained via RiboToolkit but was plotted using GraphPad Prism (GraphPad Software, Inc.).

#### Read mapping

Firstly, the processed data was mapped to Arabidopsis ncRNA (rRNA, snoRNA, tRNA) sequences (TAIR10) using Bowtie v.1.2 (parameters: “-v 2”) (Langmead et al. 2009). The ncRNA-unaligned reads were then mapped to Arabidopsis reference genome (TAIR10) using STAR v.2.5 (parameters: “ --outFilterMismatchNmax 2 --outFilterMultimapNmax 1”) (109) to yield uniquely-mapped reads. The ncRNA-unaligned reads were also mapped to RCNMV RNA1 and RNA2 using Bowtie v1.2 (parameters: “-v 2 -m 1”) (110) to yield uniquely-mapped reads. The read length distribution (Supplemental Fig. S2A; all the samples) was determined using SAMtools (111, 112). Subsequently, size filtering was done to retain only the 26-29-nt reads in the Ribo-seq alignment bam file using the reformat function in the BBMap tool (113). No size filtering was done for RNA-seq reads. The metagene analysis plots (Supplemental Fig. S3) were generated using RiboTaper (108). From the STAR-aligned files, all the RNA-seq reads were counted (parameters: “-t exon -s 0 -g gene_id”) while for Ribo-seq reads, only those mapping to the CDS were counted (parameters: “-t CDS -s 0 -g gene_id”) using the featureCounts (114). The read statistics, such as the number and proportion of raw, processed, and aligned reads can be found in Supplemental File S1.

#### Statistical analysis

Using the DESeq2 package (58), the regularized-log-transformed data from the Ribo-seq and RNA-seq counts were used for the principal component analysis (PCA). PCA showed that the major variation (PC1) among all of our datasets can be attributed to the type of library preparation, i.e., ribo-seq or RNA-seq (Supplemental Fig. S4E). The second major variation (PC2) can be attributed to the treatment, with 8 dpi RCNMV-inoculated samples forming one cluster, 5 dpi RCNMV-inoculated samples forming another cluster, and mock-inoculated samples, both 5 and 8 dpi, aggregating into a single cluster, as expected. For making correlation plots, corrplot package (115) was used only with the genes whose sum of read counts was greater than zero across all the Ribo-seq or RNA-seq samples. Pearson correlation analysis showed very high reproducibility among the biological replicates of each treatment for both Ribo-seq (Supplemental Fig. S4F) and RNA-seq dataset (Supplemental Fig. S4G). Overall, these analyses demonstrate the high-quality NGS data characteristics. DESeq2 package (58) was also used to identify the differentially expressed genes (DEGs) from the RNA-seq data and the differentially translated genes (DTGs) from the Ribo-seq data. For each of the Ribo-seq or RNA-seq data, separately, only the genes whose sum of read counts was greater than 100 across all the samples were used for analysis. All the comparisons refer to RCNMV-infected vs mock samples. Genes with absolute log2 fold change (RNA abundance) >1 and adjusted p-value < 0.05 were considered as Differentially Expressed Genes (DEGs). Genes with absolute log2 fold change (RPF abundance) >1 and adjusted p-value < 0.05 were considered as Differentially Translated Genes (DTGs). DTGs are not necessarily translationally-regulated. Genes can be identified as DTGs because of significant change in RPF abundance owing to changes in mRNA expression and/or translation efficiency. The list of DEGs and DTGs can be found in Supplemental File S2. For Fig. 2B, log_2_ fold change (RPF abundance) against log_2_ fold change (RNA abundance) was plot using only the genes that had a valid output (Non-NA) from DESeq2 (Supplemental File S4). The blue-colored genes (only-identified as DTGs) within the yellow-shaded region were considered as translationally-regulated and used for protein-domain enrichment analysis via ThaleMine (59).

#### KEGG pathway enrichment analysis

Genes were filtered based on read counts, followed by Deseq2 analysis (5 dpi RCNMV vs Mock Riboseq, 8 dpi RCNMV vs Mock Riboseq, 5 dpi RCNMV vs Mock RNAseq, 8 dpi RCNMV vs Mock RNAseq). Subsequently, we used the DEGs and DTGs from this analysis for KEGG. For background, we used the default option, we used the default option for each sample. For each of the Ribo-seq and RNA-seq datasets, the upregulated and downregulated genes were used together for the enrichment analysis. Firstly, the amino acid sequence for the DEGs/DTGs were downloaded from (Arabidopsis.org/tools/bulk/sequence) with the following parameters: dataset “Araport 11 protein sequences”, search against “get one sequence per locus (representative gene model/splice form only)”. Subsequently, KOBAS (kobas.cbi.pku.edu.cn/kobas3/genelist/) (116) was used with the following parameters: species “Arabidopsis thaliana”, input type “fasta protein sequence”, pathway “KEGG”, statistical method “hypergeometric/Fisher’s exact test”, FDR correction method “Benjamin and Hochberg (1995)”, background database “default”. Pathways with FDR < 0.05 were considered as enriched (Supplemental file S3). For Fig. 3B, log_2_ fold change (RPF abundance) against log_2_ fold change (RNA abundance) was plot using the DEGs or DTGs that were present in the enriched KEGG pathways (Supplemental file S5). There were a few DEGs that did not have a valid output (non-NA) from DESeq2 analysis of Ribo-seq data and hence, were assigned log_2_ fold change (RPF abundance) = 0 (green triangles). There were a few DTGs that did not have a valid output (non-NA) from DESeq2 analysis of RNA-seq data and hence, were assigned log_2_ fold change (RNA abundance) = 0 (blue triangles).

#### RCNMV analysis

For Fig. 4A, proportion of reads that mapped to RCNMV genome with respect to total number of Arabidopsis- and RCNMV-mapped reads was calculated (Supplemental file S1). For Fig. 4B, RCNMV read counts were included in the count table from Arabidopsis-mapped reads and input into DESeq2 for normalization. The normalized abundance was transformed via log10 (x +1) transformation. Subsequently, arithmetic mean of log10 values were calculated (equivalent to geometric mean of read counts) and plotted as a scatterplot. See Supplemental file S6 for more details. For Fig. 7, whole length of the RCNMV positive- and negative-strand-mapped RNA-seq reads (combined across all RCNMV-infected samples) were plotted over RCNMV RNA1 and RNA2 sequence. For Figs. 6 and 7, riboSeqR tool was used to visualize the coverage of RNA-seq reads (gray bars) and the Ribo-seq reads (colored bars). Here, only the position of the 5’ nucleotide of each read was plotted over the RCNMV RNA1 and RNA2 sequences.

## Data availability

The original ribo-seq data presented in the study are publicly available here: NCBI BioProject accession number PRJNA950066, https://dataview.ncbi.nlm.nih.gov/object/PRJNA950066.

## Acknowledgments

The authors thank Dr. Akshay Yadav and Dr. Gaurav Kandoi for their frequent help with bioinformatics, and Dr. Viraj Muthye for assistance with plant cultivation. The authors would also like to thank Dr. Prakrit Chotewutmontri and Dr. Polly Hsu for their advice on optimizing the Ribo-seq protocol. F.L was supported by BBSRC grants (BB/X001261/1 and BB/V006096/1) to BYWC. This project was funded by Iowa State University Plant Sciences Institute Faculty Scholar award to WAM. This paper of the Iowa Agriculture and Home Economics Experiment Station, Ames, IA, Project No. 3808 was supported in part by Hatch Act and State of Iowa funds.

## Conflict of Interest

The authors declare no conflict of interest associated with the work described in this manuscript.

## Notes

https://dataview.ncbi.nlm.nih.gov/object/PRJNA950066

### Competing Interest Statement

The authors have declared no competing interest.

### Summary of Updates

1. Removed revision date for journal submission. 2. Made ribo-seq consistently lower case. 3. Submitted figures as higher resolutioin .tif files. 3. Added 4 supplemental figures.

## References

1. Collum TD, Padmanabhan MS, Hsieh YC, Culver JN. 2016. Tobacco mosaic virus-directed reprogramming of auxin/indole acetic acid protein transcriptional responses enhances virus phloem loading. Proc Natl Acad Sci U S A 113:E2740–9.

2. Nicaise V, Candresse T. 2017. Plum pox virus capsid protein suppresses plant pathogen-associated molecular pattern (PAMP)-triggered immunity. Mol Plant Pathol 18:878–886.

3. Zhang X, Hong H, Yan J, Yuan Y, Feng M, Liu Q, Zhao Y, Yang T, Huang S, Wang C, Zhao R, Zuo W, Liu S, Ding Z, Huang C, Zhang Z, Kundu JK, Tao X. 2024. Autophagy plays an antiviral defence role against tomato spotted wilt orthotospovirus and is counteracted by viral effector NSs. Mol Plant Pathol 25:e70012.

4. Jiang T, Hao T, Chen W, Li C, Pang S, Fu C, Cheng J, Zhang C, Ghorbanpour M, Miao S. 2025. Reprogrammed Plant Metabolism During Viral Infections: Mechanisms, Pathways and Implications. Mol Plant Pathol 26:e70066.

5. Bengyella L, Waikhom SD, Allie F, Rey C. 2015. Virus tolerance and recovery from viral induced-symptoms in plants are associated with transcriptome reprograming. Plant Mol Biol 89:243–52.

6. Garcia-Ruiz H, Szurek B, Van den Ackerveken G. 2021. Stop helping pathogens: engineering plant susceptibility genes for durable resistance. Curr Opin Biotechnol 70:187–195.

7. Gal-On A, Fuchs M, Gray S. 2017. Generation of novel resistance genes using mutation and targeted gene editing. Curr Opin Virol 26:98–103.

8. Medzihradszky A, Gyula P, Sos-Hegedus A, Szittya G, Burgyan J. 2019. Transcriptome reprogramming in the shoot apical meristem of CymRSV-infected Nicotiana benthamiana plants associates with viral exclusion and the lack of recovery. Mol Plant Pathol 20:1748–1758.

9. Herath V, Casteel CL, Verchot J. 2025. Comprehensive transcriptomic analysis reveals turnip mosaic virus infection and its aphid vector Myzus persicae cause large changes in gene regulatory networks and co-transcription of alternative spliced mRNAs in Arabidopsis thaliana. BMC Plant Biol 25:128.

10. Samarskaya V, Spechenkova N, Kalinina NO, Love AJ, Taliansky M. 2025. The Emerging Role of Omics-Based Approaches in Plant Virology. Viruses 17.

11. Bhattacharjee S, Zamora A, Azhar MT, Sacco MA, Lambert LH, Moffett P. 2009. Virus resistance induced by NB-LRR proteins involves Argonaute4-dependent translational control. Plant J 58:940–51.

12. Zorzatto C, Machado JP, Lopes KV, Nascimento KJ, Pereira WA, Brustolini OJ, Reis PA, Calil IP, Deguchi M, Sachetto-Martins G, Gouveia BC, Loriato VA, Silva MA, Silva FF, Santos AA, Chory J, Fontes EP. 2015. NIK1-mediated translation suppression functions as a plant antiviral immunity mechanism. Nature 520:679–82.

13. Machado JPB, Calil IP, Santos AA, Fontes EPB. 2017. Translational control in plant antiviral immunity. Genet Mol Biol 40:292–304.

14. Miller WA, Shen R, Staplin W, Kanodia P. 2016. Noncoding RNAs of plant viruses and viroids: Sponges of host translation and RNA interference machinery. Mol Plant Microbe Interact 29:156–164.

15. Wu H, Qu X, Dong Z, Luo L, Shao C, Forner J, Lohmann JU, Su M, Xu M, Liu X, Zhu L, Zeng J, Liu S, Tian Z, Zhao Z. 2020. WUSCHEL triggers innate antiviral immunity in 14. plant stem cells. Science 370:227–231.

16. Moeller JR, Moscou MJ, Bancroft T, Skadsen RW, Wise RP, Whitham SA. 2012. Differential accumulation of host mRNAs on polyribosomes during obligate pathogen-plant interactions. Mol Biosyst 8:2153–65.

17. Wang D, Maule AJ. 1995. Inhibition of host gene expression associated with plant virus replication. Science 267:229–231.

18. Eskelin K, Varjosalo M, Ravantti J, Makinen K. 2019. Ribosome profiles and riboproteomes of healthy and Potato virus A- and Agrobacterium-infected Nicotiana benthamiana plants. Mol Plant Pathol 20:392–409.

19. Collum TD, Stone AL, Sherman DJ, Rogers EE, Dardick C, Culver JN. 2020. Translatome Profiling of Plum Pox Virus-Infected Leaves in European Plum Reveals Temporal and Spatial Coordination of Defense Responses in Phloem Tissues. Mol Plant Microbe Interact 33:66–77.

20. Ingolia NT, Ghaemmaghami S, Newman JR, Weissman JS. 2009. Genome-wide analysis in vivo of translation with nucleotide resolution using ribosome profiling. Science 324:218–223.

21. Stern-Ginossar N, Ingolia NT. 2015. Ribosome Profiling as a Tool to Decipher Viral Complexity. Annu Rev Virol 2:335–49.

22. Irigoyen N, Firth AE, Jones JD, Chung BY, Siddell SG, Brierley I. 2016. High-Resolution Analysis of Coronavirus Gene Expression by RNA Sequencing and Ribosome Profiling. PLoS Pathog 12:e1005473.

23. Irigoyen N, Dinan AM, Brierley I, Firth AE. 2018. Ribosome profiling of the retrovirus murine leukemia virus. Retrovirology 15:10.

24. Pallares HM, Gonzalez Lopez Ledesma MM, Oviedo-Rouco S, Castellano LA, Costa Navarro GS, Fernandez-Alvarez AJ, D’Andreiz MJ, Aldas-Bulos VD, Alvarez DE, Bazzini AA, Gamarnik AV. 2024. Zika virus non-coding RNAs antagonize antiviral responses by PKR-mediated translational arrest. Nucleic Acids Res doi:10.1093/nar/gkae507.

25. Cook GM, Brown K, Shang P, Li Y, Soday L, Dinan AM, Tumescheit C, Mockett APA, Fang Y, Firth AE, Brierley I. 2022. Ribosome profiling of porcine reproductive and respiratory syndrome virus reveals novel features of viral gene expression. Elife 11.

26. Zhao J, Huang Y, Liukang C, Yang R, Tang L, Sun L, Zhao Y, Zhang G. 2024. Dissecting infectious bronchitis virus-induced host shutoff at the translation level. J Virol 98:e0083024.

27. Castellano LA, McNamara RJ, Pallares HM, Gamarnik AV, Alvarez DE, Bazzini AA. 2024. Dengue virus preferentially uses human and mosquito non-optimal codons. Mol Syst Biol 20:1085–1108.

28. Wu HL, Jen J, Hsu PY. 2024. What, where, and how: Regulation of translation and the translational landscape in plants. Plant Cell 36:1540–1564.

29. Kawamoto N, Iwasaki S. 2026. Translating the green code: Ribo-Seq for photosynthetic eukaryotes in focus. J Biochem doi:10.1093/jb/mvag004.

30. Bai B, Qi R, Song W, Nijveen H, Bentsink L. 2026. Translational landscape during seed germination revealed by ribosome profiling. Plant J 125:e70663.

31. Xu T, Lei L, Shi J, Wang X, Chen J, Xue M, Sun S, Zhan B, Xia Z, Jiang N, Zhou T, Lai J, Fan Z. 2019. Characterization of maize translational responses to sugarcane mosaic virus infection. Virus Res 259:97–107.

32. Chiu C-W, Li Y-R, Lin C-Y, Yeh H-H, Liu M-J. 2022. Translation initiation landscape profiling reveals hidden open-reading frames required for the pathogenesis of tomato yellow leaf curl Thailand virus. The Plant Cell 34:1804–1821.

33. Lukhovitskaya N, Brown K, Hua L, Pate AE, Carr JP, Firth AE. 2024. A novel ilarvirus protein CP-RT is expressed via stop codon readthrough and suppresses RDR6-dependent RNA silencing. PLoS Pathog 20:e1012034.

34. Gong P, Gao M, Chen Y, Zhang M, Huang Y, Hu X, Zhao S, Zhang H, Pan M, Cao B, Shen Q, Liu Y, Lozano-Duran R, Wang A, Zhou X, Li F. 2025. Cucumber green mottle mosaic virus encodes additional small proteins with specific subcellular localizations and virulence function. Sci China Life Sci 68:1815–1827.

35. Wang C, Tang Y, Zhou C, Li S, Chen J, Sun Z. 2024. RNA-seq and Ribosome Profiling Reveal the Translational Landscape of Rice in Response to Rice Stripe Virus Infection. Viruses 16.

36. Gould AR, Francki RI, Hatta T, Hollings M. 1981. The bipartite genome of red clover necrotic mosaic virus. Virology 108:499–506.

37. Okuno T, Hiruki C. 2013. Molecular biology and epidemiology of dianthoviruses. Adv Virus Res 87:37–74.

38. Xiong Z, Lommel SA. 1989. The complete nucleotide sequence and genome organization of red clover necrotic mosaic virus RNA-1. Virology 171:543–554.

39. Kim KH, Lommel SA. 1994. Identification and Analysis of the Site of -1 Ribosomal Frameshifting in Red Clover Necrotic Mosaic Virus. Virology 200:574–582.

40. Xiong Z, Kim KH, Giesman-Cookmeyer D, Lommel SA. 1993. The roles of the red clover necrotic mosaic virus capsid and cell-to-cell movement proteins in systemic infection. Virology 192:27–32.

41. Sit TL, Vaewhongs AA, Lommel SA. 1998. RNA-mediated transactivation of transcription from a viral RNA. Science 281:829–832.

42. Lommel SA, Weston-Fina M, Xiong Z, Lomonossoff GP. 1988. The nucleotide sequence and gene organization of red clover necrotic mosaic virus RNA-2. Nucleic Acids Res 16:8587–602.

43. Iwakawa HO, Mizumoto H, Nagano H, Imoto Y, Takigawa K, Sarawaneeyaruk S, Kaido M, Mise K, Okuno T. 2008. A viral noncoding RNA generated by cis-element-mediated protection against 5’->3’ RNA decay represses both cap-independent and cap-dependent translation. J Virol 82:10162–74.

44. Steckelberg AL, Akiyama BM, Costantino DA, Sit TL, Nix JC, Kieft JS. 2018. A folded viral noncoding RNA blocks host cell exoribonucleases through a conformationally dynamic RNA structure. Proc Natl Acad Sci U S A 115:6404–6409.

45. Mizumoto H, Tatsuta M, Kaido M, Mise K, Okuno T. 2003. Cap-independent translational enhancement by the 3’ untranslated region of red clover necrotic mosaic virus RNA1. J Virol 77:12113–12121.

46. Kraft JJ, Treder K, Peterson MS, Miller WA. 2013. Cation-dependent folding of 3’ cap-independent translation elements facilitates interaction of a 17-nucleotide conserved sequence with eIF4G. Nucleic Acids Res 41:3398–413.

47. Treder K, Pettit Kneller EL, Allen EM, Wang Z, Browning KS, Miller WA. 2008. The 3’ cap-independent translation element of Barley yellow dwarf virus binds eIF4F via the eIF4G subunit to initiate translation. RNA 14:134–147.

48. Zhao P, Liu Q, Miller WA, Goss DJ. 2017. Eukaryotic translation initiation factor 4G (eIF4G) coordinates interactions with eIF4A, eIF4B and eIF4E in binding and translation of the barley yellow dwarf virus 3’ cap-independent translation element (BTE). J Biol Chem 292:5921–5931.

49. Li Z, Pogany J, Tupman S, Esposito AM, Kinzy TG, Nagy PD. 2010. Translation elongation factor 1A facilitates the assembly of the tombusvirus replicase and stimulates minus-strand synthesis. PLoS pathogens 6:e1001175.

50. Pogany J, Nagy PD. 2015. Activation of Tomato Bushy Stunt Virus RNA-Dependent RNA Polymerase by Cellular Heat Shock Protein 70 Is Enhanced by Phospholipids In Vitro. J Virol 89:5714–23.

51. Kovalev N, Pogany J, Nagy PD. 2020. Reconstitution of an RNA Virus Replicase in Artificial Giant Unilamellar Vesicles Supports Full Replication and Provides Protection for the Double-Stranded RNA Replication Intermediate. J Virol 94.

52. Nagy PD. 2016. Tombusvirus-Host Interactions: Co-Opted Evolutionarily Conserved Host Factors Take Center Court. Annu Rev Virol 3:491–515.

53. Chung BY, Hardcastle TJ, Jones JD, Irigoyen N, Firth AE, Baulcombe DC, Brierley I. 2015. The use of duplex-specific nuclease in ribosome profiling and a user-friendly software package for Ribo-seq data analysis. RNA 21:1731–45.

54. Kanodia P, Miller WA. 2022. Effects of the Noncoding Subgenomic RNA of Red Clover Necrotic Mosaic Virus in Virus Infection. J Virol 96:e0181521.

55. Deleris A, Gallego-Bartolome J, Bao J, Kasschau KD, Carrington JC, Voinnet O. 2006. Hierarchical action and inhibition of plant Dicer-like proteins in antiviral defense. Science 313:68–71.

56. Xie Z, Allen E, Wilken A, Carrington JC. 2005. DICER-LIKE 4 functions in trans-acting small interfering RNA biogenesis and vegetative phase change in Arabidopsis thaliana. Proc Natl Acad Sci U S A 102:12984–9.

57. Hsu PY, Calviello L, Wu HL, Li FW, Rothfels CJ, Ohler U, Benfey PN. 2016. Super-resolution ribosome profiling reveals unannotated translation events in Arabidopsis. Proc Natl Acad Sci U S A doi:10.1073/pnas.1614788113.

58. Love MI, Huber W, Anders S. 2014. Moderated estimation of fold change and dispersion for RNA-seq data with DESeq2. Genome Biol 15:550.

59. Krishnakumar V, Contrino S, Cheng CY, Belyaeva I, Ferlanti ES, Miller JR, Vaughn MW, Micklem G, Town CD, Chan AP. 2017. ThaleMine: A Warehouse for Arabidopsis Data Integration and Discovery. Plant Cell Physiol 58:e4.

60. Macho AP, Lozano-Duran R. 2019. Molecular dialogues between viruses and receptor-like kinases in plants. Mol Plant Pathol 20:1191–1195.

61. Yu TY, Sun MK, Liang LK. 2021. Receptors in the Induction of the Plant Innate Immunity. Mol Plant Microbe Interact 34:587–601.

62. Snoeck S, Johanndrees O, Nurnberger T, Zipfel C. 2025. Plant pattern recognition receptors: from evolutionary insight to engineering. Nat Rev Genet 26:268–278.

63. Deng Y, Srivastava R, Quilichini TD, Dong H, Bao Y, Horner HT, Howell SH. 2016. IRE1, a component of the unfolded protein response signaling pathway, protects pollen development in Arabidopsis from heat stress. Plant J 88:193–204.

64. Howell SH. 2013. Endoplasmic reticulum stress responses in plants. Annu Rev Plant Biol 64:477–99.

65. Adhikari B, Gayral M, Herath V, Bedsole CO, Kumar S, Ball H, Atallah O, Shaw B, Pajerowska-Mukhtar KM, Verchot J. 2024. bZIP60 and Bax inhibitor 1 contribute IRE1- dependent and independent roles to potexvirus infection. New Phytol 243:1172–1189.

66. Chotewutmontri P, Barkan A. 2016. Dynamics of Chloroplast Translation during Chloroplast Differentiation in Maize. PLoS Genet 12:e1006106.

67. Guenther RH, Sit TL, Gracz HS, Dolan MA, Townsend HL, Liu G, Newman WH, Agris PF, Lommel SA. 2004. Structural characterization of an intermolecular RNA-RNA interaction involved in the transcription regulation element of a bipartite plant virus. Nucleic Acids Res 32:2819–28.

68. Chung BY, Hardcastle TJ, Jones JD, Irigoyen N, Firth AE, Baulcombe DC, Brierley I. 2015. The use of duplex-specific nuclease in ribosome profiling and a user-friendly software package for Ribo-seq data analysis. RNA 21:1731–1745.

69. López-Márquez D, Del-Espino Á, Ruiz-Albert J, Bejarano ER, Brodersen P, Beuzón CR. 2023. Regulation of plant immunity via small RNA-mediated control of NLR expression. Journal of Experimental Botany 74:6052–6068.

70. Nielsen CPS, Arribas-Hernández L, Han L, Reichel M, Woessmann J, Daucke R, Bressendorff S, López-Márquez D, Andersen SU, Pumplin N, Schoof EM, Brodersen P. 2024. Evidence for an RNAi-independent role of Arabidopsis DICER-LIKE2 in growth inhibition and basal antiviral resistance. The Plant Cell 36:2289–2309.

71. Lewsey MG, Carr JP. 2009. Effects of DICER-like proteins 2, 3 and 4 on cucumber mosaic virus and tobacco mosaic virus infections in salicylic acid-treated plants. J Gen Virol 90:3010–3014.

72. Ju C, Li L, Zhang Y, Meng Y, Liu W. 2025. Tumor necrosis factor receptor-associated factors (TRAFs): Key regulatory factors in antiviral defence. Microb Pathog 206:107755.

73. Qi H, Xia FN, Xiao S, Li J. 2022. TRAF proteins as key regulators of plant development and stress responses. J Integr Plant Biol 64:431–448.

74. Huang S, Chen X, Zhong X, Li M, Ao K, Huang J, Li X. 2016. Plant TRAF Proteins Regulate NLR Immune Receptor Turnover. Cell Host Microbe 19:204–15.

75. Adhikari B, Verchot J, Brandizzi F, Ko DK. 2025. ER stress and viral defense: Advances and future perspectives on plant unfolded protein response in pathogenesis. J Biol Chem 301:108354.

76. Turner KA, Sit TL, Callaway AS, Allen NS, Lommel SA. 2004. Red clover necrotic mosaic virus replication proteins accumulate at the endoplasmic reticulum. Virology 320:276–90.

77. Kusumanegara K, Mine A, Hyodo K, Kaido M, Mise K, Okuno T. 2012. Identification of domains in p27 auxiliary replicase protein essential for its association with the endoplasmic reticulum membranes in Red clover necrotic mosaic virus. Virology 433:131–41.

78. Kanodia P, Vijayapalani P, Srivastava R, Bi R, Liu P, Miller WA, Howell SH. 2020. Control of translation during the unfolded protein response in maize seedlings: Life without PERKs. Plant Direct 4:e00241.

79. May J, Johnson P, Saleem H, Simon AE. 2017. A Sequence-Independent, Unstructured IRES is Responsible for Internal Expression of the Coat Protein of Turnip Crinkle Virus. J Virol doi:10.1128/jvi.02421-16.

80. Carino EJ, Scheets K, Miller WA. 2020. The RNA of Maize Chlorotic Mottle Virus, an Obligatory Component of Maize Lethal Necrosis Disease, Is Translated via a Variant Panicum Mosaic Virus-Like Cap-Independent Translation Element. J Virol 94.

81. Barry JK, Miller WA. 2002. A -1 ribosomal frameshift element that requires base pairing across four kilobases suggests a mechanism of regulating ribosome and replicase traffic on a viral RNA. Proc Natl Acad Sci U S A 99:11133–11138.

82. Tamm T, Suurvali J, Lucchesi J, Olspert A, Truve E. 2009. Stem-loop structure of Cocksfoot mottle virus RNA is indispensable for programmed -1 ribosomal frameshifting. Virus Res 146:73–80.

83. Tajima Y, Iwakawa HO, Kaido M, Mise K, Okuno T. 2011. A long-distance RNA-RNA interaction plays an important role in programmed -1 ribosomal frameshifting in the translation of p88 replicase protein of Red clover necrotic mosaic virus. Virology 417:169–178.

84. Kim KH, Lommel SA. 1998. Sequence element required for efficient -1 ribosomal frameshifting in red clover necrotic mosaic dianthovirus. Virology 250:50–9.

85. Brierley I, Gilbert RJC, Pennell S. 2010. Pseudoknot-dependent programmed -1 ribosomal frameshifting: Structures, mechanisms and models, p 149-174. In Atkins JF, Gesteland RF (ed), Recoding: Expansion of Decoding Rules Enriches Gene Expression. Springer, London.

86. Dinman JD. 2012. Mechanisms and implications of programmed translational frameshifting. Wiley Interdiscip Rev RNA 3:661–73.

87. Kibe A, Buck S, Gribling-Burrer AS, Gilmer O, Bohn P, Koch T, Mireisz CN, Schlosser A, Erhard F, Smyth RP, Caliskan N. 2025. The translational landscape of HIV-1 infected cells reveals key gene regulatory principles. Nat Struct Mol Biol 32:841–852.

88. Finkel Y, Gluck A, Nachshon A, Winkler R, Fisher T, Rozman B, Mizrahi O, Lubelsky Y, Zuckerman B, Slobodin B, Yahalom-Ronen Y, Tamir H, Ulitsky I, Israely T, Paran N, Schwartz M, Stern-Ginossar N. 2021. SARS-CoV-2 uses a multipronged strategy to impede host protein synthesis. Nature 594:240–245.

89. Napthine S, Ling R, Finch LK, Jones JD, Bell S, Brierley I, Firth AE. 2017. Protein-directed ribosomal frameshifting temporally regulates gene expression. Nat Commun 8:15582.

90. Kontos H, Napthine S, Brierley I. 2001. Ribosomal pausing at a frameshifter RNA pseudoknot is sensitive to reading phase but shows little correlation with frameshift efficiency. Mol Cell Biol 21:8657–8670.

91. Bhatt PR, Scaiola A, Loughran G, Leibundgut M, Kratzel A, Meurs R, Dreos R, O’Connor KM, McMillan A, Bode JW, Thiel V, Gatfield D, Atkins JF, Ban N. 2021. Structural basis of ribosomal frameshifting during translation of the SARS-CoV-2 RNA genome. Science 372:1306–1313.

92. Yanagitani K, Kimata Y, Kadokura H, Kohno K. 2011. Translational pausing ensures membrane targeting and cytoplasmic splicing of XBP1u mRNA. Science 331:586–9.

93. Cao J, Geballe AP. 1995. Translational inhibition by a human cytomegalovirus upstream open reading frame despite inefficient utilization of its AUG codon. Journal of Virology 69:1030–1036.

94. Melnikov S, Mailliot J, Rigger L, Neuner S, Shin BS, Yusupova G, Dever TE, Micura R, Yusupov M. 2016. Molecular insights into protein synthesis with proline residues. EMBO reports 17:1776–1784.

95. Panteleev PV, Pichkur EB, Kruglikov RN, Paleskava A, Shulenina OV, Bolosov IA, Bogdanov IV, Safronova VN, Balandin SV, Marina VI, Kombarova TI, Korobova OV, Shamova OV, Myasnikov AG, Borzilov AI, Osterman IA, Sergiev PV, Bogdanov AA, Dontsova OA, Konevega AL, Ovchinnikova TV. 2024. Rumicidins are a family of mammalian host-defense peptides plugging the 70S ribosome exit tunnel. Nature Communications 15:8925.

96. Wruck F, Katranidis A, Nierhaus KH, Büldt G, Hegner M. 2017. Translation and folding of single proteins in real time. Proceedings of the National Academy of Sciences 114:E4399–E4407.

97. Simms CL, Yan LL, Zaher HS. 2017. Ribosome Collision Is Critical for Quality Control during No-Go Decay. Mol Cell 68:361–373 e5.

98. Navickas A, Chamois S, Saint-Fort R, Henri J, Torchet C, Benard L. 2020. No-Go Decay mRNA cleavage in the ribosome exit tunnel produces 5’-OH ends phosphorylated by Trl1. Nat Commun 11:122.

99. Tatsuta M, Mizumoto H, Kaido M, Mise K, Okuno T. 2005. The red clover necrotic mosaic virus RNA2 trans-activator is also a cis-acting RNA2 replication element. J Virol 79:978–86.

100. den Boon JA, Ahlquist P. 2010. Organelle-like membrane compartmentalization of positive-strand RNA virus replication factories. Annual review of microbiology 64:241–56.

101. Mizumoto H, Iwakawa HO, Kaido M, Mise K, Okuno T. 2006. Cap-independent translation mechanism of red clover necrotic mosaic virus RNA2 differs from that of RNA1 and is linked to RNA replication. J Virol 80:3781–3791.

102. Kanodia P, Prasanth KR, Roa-Linares VC, Bradrick SS, Garcia-Blanco MA, Miller WA. 2020. A rapid and simple quantitative method for specific detection of smaller coterminal RNA by PCR (DeSCo-PCR): application to the detection of viral subgenomic RNAs. RNA 26:888–901.

103. Boyes DC, Zayed AM, Ascenzi R, McCaskill AJ, Hoffman NE, Davis KR, Gorlach J. 2001. Growth stage-based phenotypic analysis of Arabidopsis: a model for high throughput functional genomics in plants. Plant Cell 13:1499–510.

104. Chung BYW, Balcerowicz M, Di Antonio M, Jaeger KE, Geng F, Franaszek K, Marriott P, Brierley I, Firth AE, Wigge PA. 2020. An RNA thermoswitch regulates daytime growth in Arabidopsis. Nat Plants 6:522–532.

105. Sonia J, Kanodia P, Lozier Z, Miller WA. 2024. Ribosome Profiling of Plants. Methods Mol Biol 2724:139–163.

106. Martin M. 2015. Cutadapt removes adapter sequences from high-throughput sequencing reads. EMBnetjournal 17.

107. Liu Q, Shvarts T, Sliz P, Gregory RI. 2020. RiboToolkit: an integrated platform for analysis and annotation of ribosome profiling data to decode mRNA translation at codon resolution. Nucleic Acids Res 48:W218–W229.

108. Calviello L, Mukherjee N, Wyler E, Zauber H, Hirsekorn A, Selbach M, Landthaler M, Obermayer B, Ohler U. 2016. Detecting actively translated open reading frames in ribosome profiling data. Nat Methods 13:165–70.

109. Dobin A, Davis CA, Schlesinger F, Drenkow J, Zaleski C, Jha S, Batut P, Chaisson M, Gingeras TR. 2013. STAR: ultrafast universal RNA-seq aligner. Bioinformatics 29:15–21.

110. Langmead B, Trapnell C, Pop M, Salzberg SL. 2009. Ultrafast and memory-efficient alignment of short DNA sequences to the human genome. Genome Biol 10:R25.

111. Danecek P, Bonfield JK, Liddle J, Marshall J, Ohan V, Pollard MO, Whitwham A, Keane T, McCarthy SA, Davies RM, Li H. 2021. Twelve years of SAMtools and BCFtools. Gigascience 10.

112. Li H, Handsaker B, Wysoker A, Fennell T, Ruan J, Homer N, Marth G, Abecasis G, Durbin R, Genome Project Data Processing S. 2009. The Sequence Alignment/Map format and SAMtools. Bioinformatics 25:2078–9.

113. Bushnell B. 2014. BBMap: A Fast, Accurate, Splice-Aware Aligner, on Lawrence Berkeley National Laboratory. https://escholarship.org/uc/item/1h3515gn. Accessed

114. Liao Y, Smyth GK, Shi W. 2014. featureCounts: an efficient general purpose program for assigning sequence reads to genomic features. Bioinformatics 30:923–30.

115. Wei T, Simko V. 2021. R package “corrplot”: Visualization of a Correlation Matrix (Version 0.90). https://github.com/taiyun/corrplot. Accessed

116. Bu D, Luo H, Huo P, Wang Z, Zhang S, He Z, Wu Y, Zhao L, Liu J, Guo J, Fang S, Cao W, Yi L, Zhao Y, Kong L. 2021. KOBAS-i: intelligent prioritization and exploratory visualization of biological functions for gene enrichment analysis. Nucleic Acids Res 49:W317–W325.

