## Supplementary figures and images for "Changes in the translational landscape during red clover necrotic mosaic virus infection"

### Supplemental Fig. 4

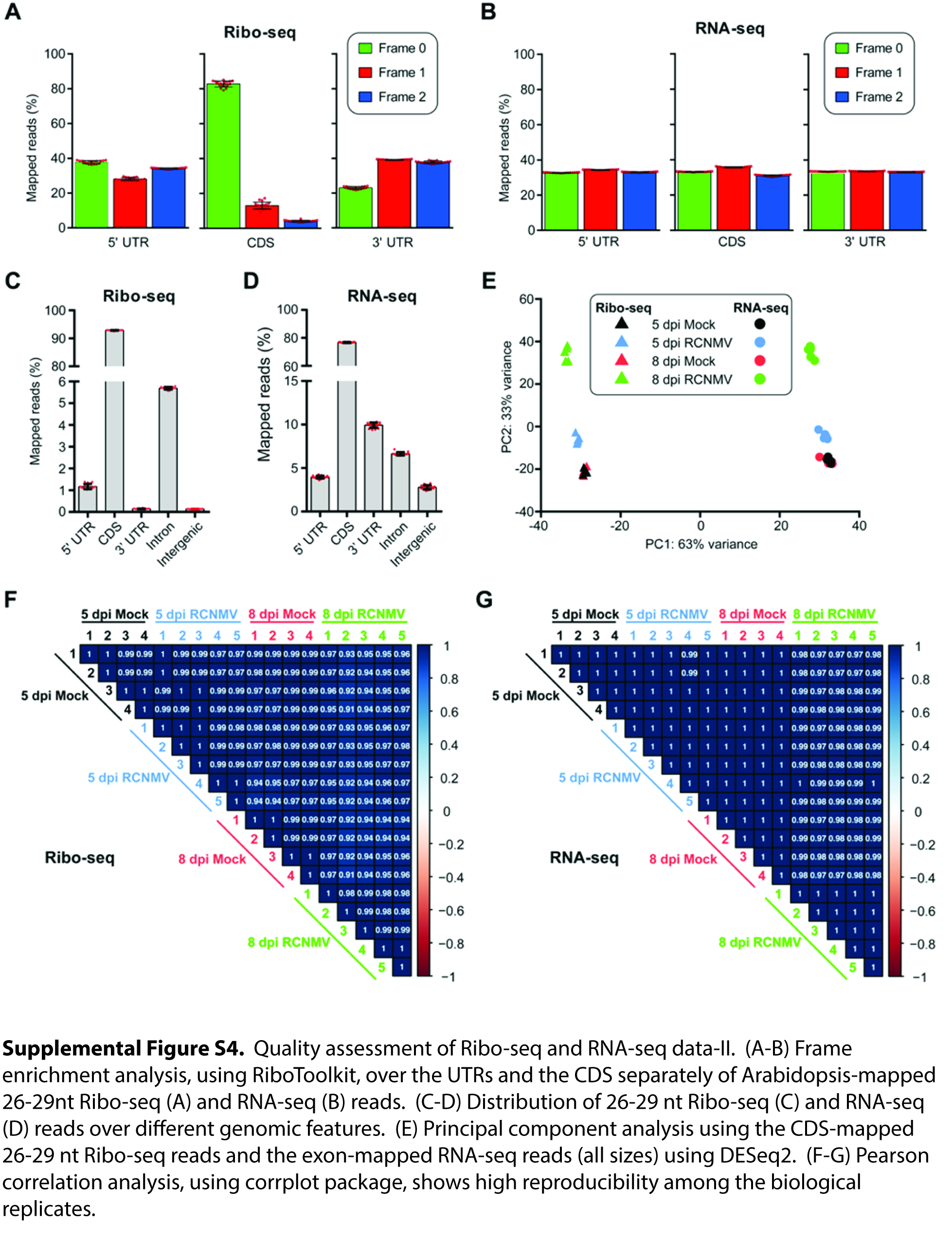

### Supplemental Fig. S1

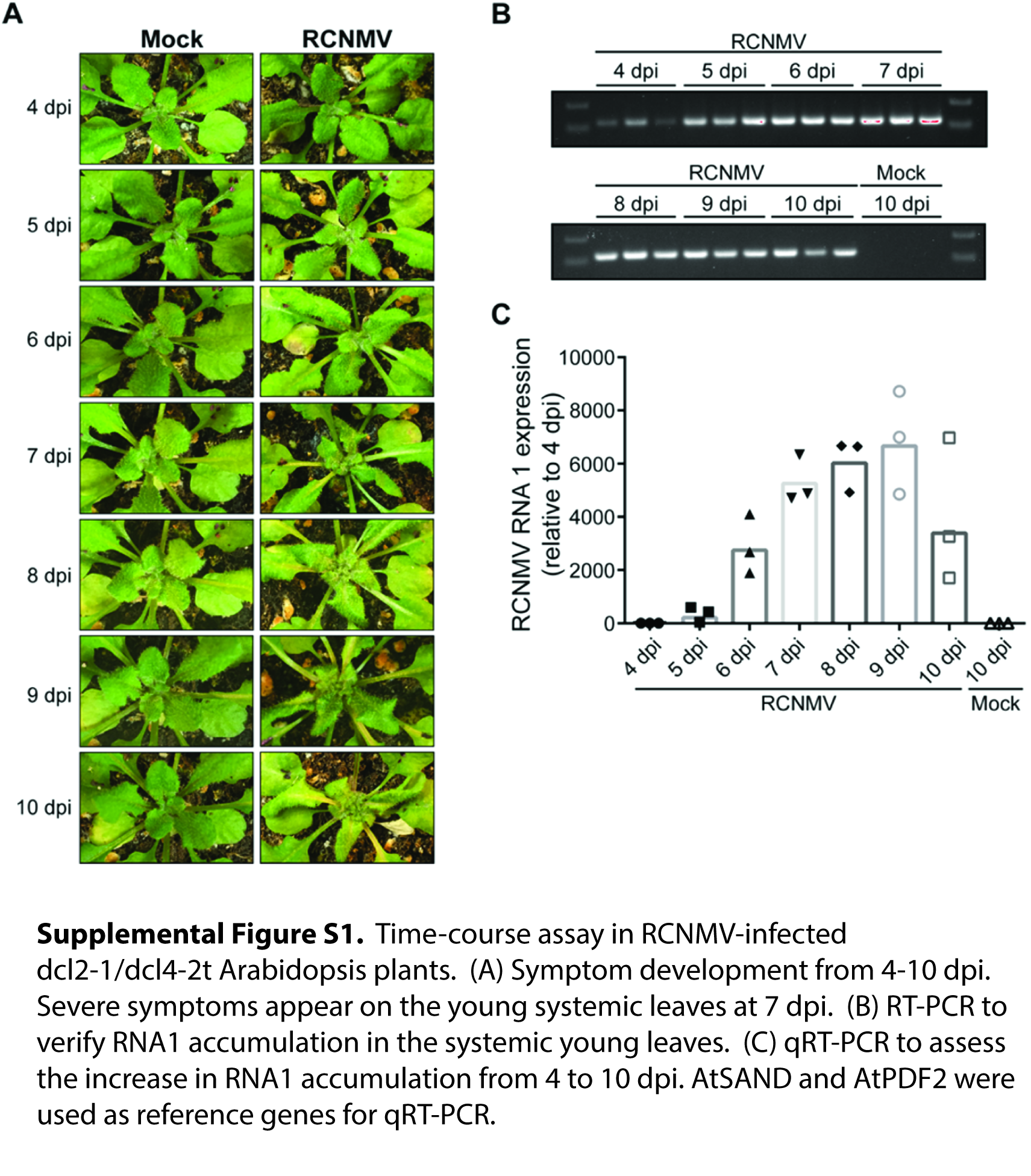

### Supplemental Fig. S2

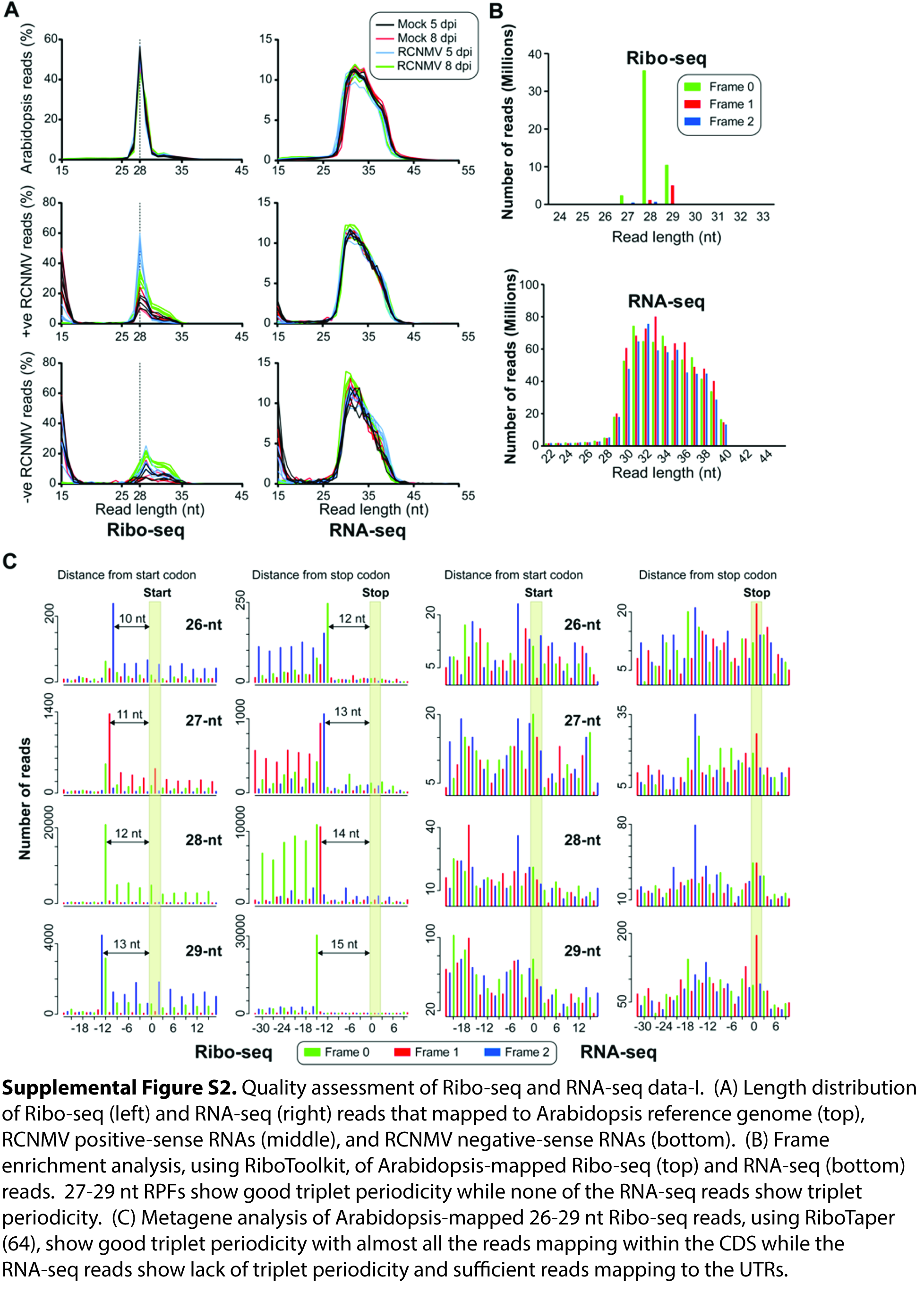

### Supplemental Fig. S3

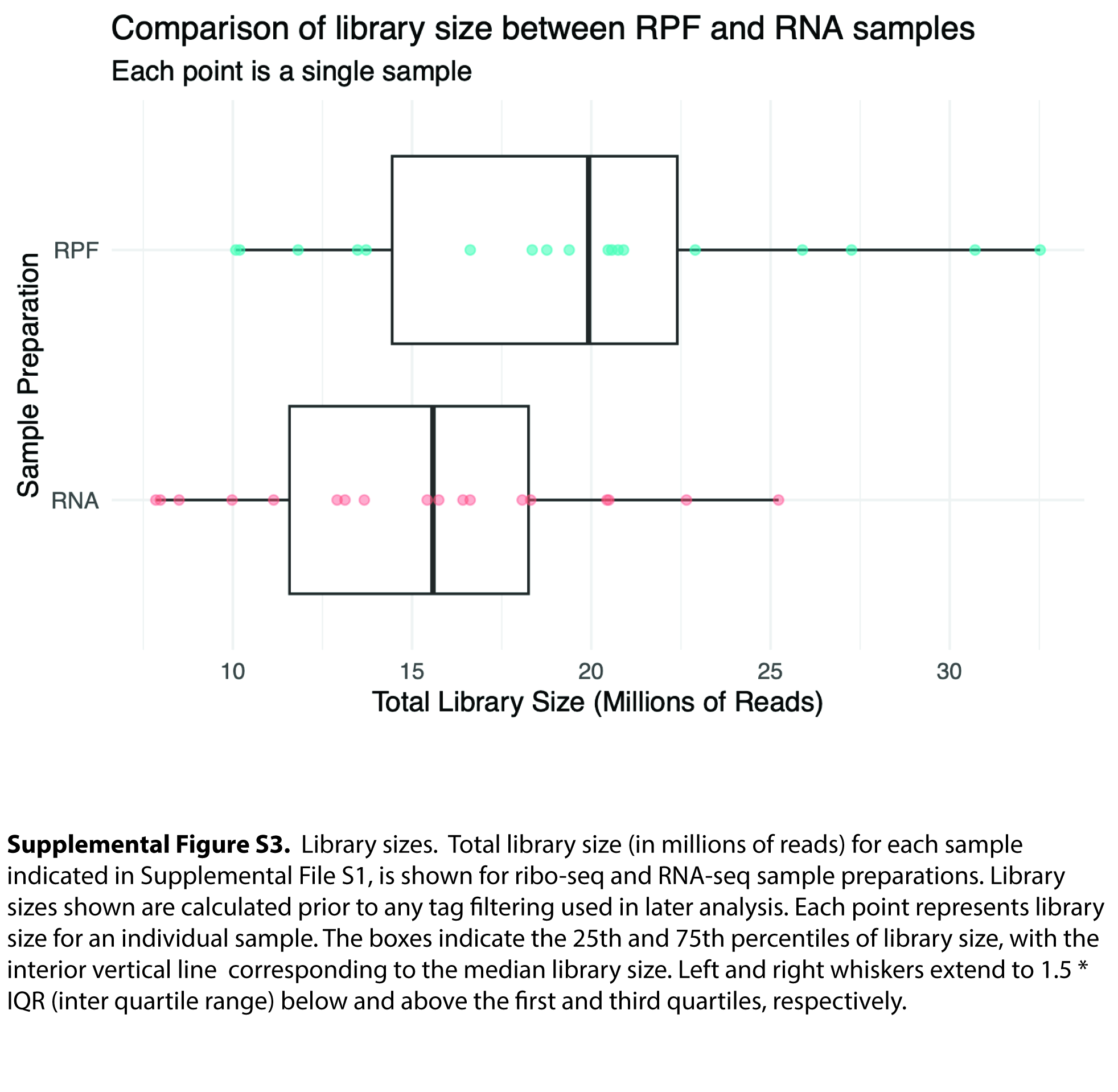
